# Effects of atmospheric CO_2_ concentration on transpiration and leaf elongation responses to drought in wheat, perennial ryegrass and tall fescue

**DOI:** 10.1101/2023.12.06.570419

**Authors:** Victoria Acker, Jean-Louis Durand, Cédric Perrot, Eric Roy, Elzbieta Frak, Romain Barillot

## Abstract

We studied the effects of atmospheric CO_2_ concentration ([CO_2_]) on the leaf growth response to drought in perennial ryegrass, tall fescue and wheat.

Plants were grown in growth chambers at either 200 or 800 ppm of CO_2_. At leaf 6-7 unfolding, half of the plants were subjected to severe drought. Leaf elongation rate (LER) was measured daily, while plant transpiration was continuously recorded gravimetrically. Water-soluble carbohydrate concentration, water and osmotic potentials in the leaf growing zone were measured at drought onset, at mid-drought and at the time of leaf growth cessation.

[CO_2_] caused stomata closure and therefore reduction of instantaneous transpiration rate and water loss. As a result, CO_2_ mitigated the impacts of drought on LER and delayed growth cessation for all three species. For ryegrass, LER and soil relative water content (SRWC) relation was improved with CO_2_, presumably due to a better stomatal regulation. CO_2_ did not affect nighttime water potential nor osmotic potential of the growing zone. Related to leaf growth, we observed the main effect of CO_2_ on tillering but no effect on the plant development. In total, water consumption was similar (wheat, tall fescue) or greater (ryegrass) with CO_2_.

## 1. INTRODUCTION

The ongoing climate change is characterised by a rapid rise in atmospheric CO_2_ concentration ([CO_2_]) and the resulting warming. As a consequence, the intensity and frequency of droughts are expected to increase (IPCC, 2023), putting current agricultural systems at risk. About 80% of the global area used by agriculture is rain-fed, but much of this area will need additional irrigation in the future, posing new challenges for agricultural practices. It is therefore crucial to improve plant adaptability and agronomic performances for future growth conditions. In this regard, plant morphogenesis and in particular leaf growth is a key process, as it is prominent for light interception, transpiration, water uptake and CO_2_ assimilation and therefore total plant productivity. As [CO_2_] increases photosynthesis and decreases transpiration, while drought reduces both processes, complex interactions are expected on leaf growth making it difficult to anticipate the effect of climate change (Kirschbaum, 2004).

With more than 10 000 species (Bouchenak-Khelladi *et al*., 2010), Poaceae represent a large part of the flora found in terrestrial ecosystems and comprise the basis of many agrosystems. For these species, leaf growth occurs at the base of the developing leaf which is enclosed in the older sheaths (Begg and Wright, 1962; Esau, 1977). As a consequence, the growing zone of the leaf (GZ) is not directly exposed to light, is not photosynthetic, does not transpire and is entirely dependent on the import of assimilates from mature tissues. Moreover, GZs are characterised by a longitudinal gradient of tissue development resulting from continuous cell production in the intercalary meristem at the leaf base, cell expansion, and differentiation (Schnyder *et al*., 1990). Cellular expansion results from a coupling between: (1) an irreversible expansion of the cell wall when turgor pressure exceeds a yield threshold, (2) an uptake of water into the expanding cell, driven by the water potential gradient between the cell and the xylem, (3) an osmotic adjustment, through an uptake of solutes, which allows the maintenance of turgor pressure and water flow and (4) metabolic processes involved in cell wall relaxation (Barlow, 1986; Cosgrove, 2018; Lockhart, 1965; Martre *et al*., 1999; Pantin *et al*., 2012; Passioura & Fry, 1992; Ray & Green, 1972).

Several mechanisms involved in leaf growth are affected by [CO_2_] and water availability. Each factor has been reported independently in many studies, but only a few investigated their interactions that affect both plant photosynthetic activity and water status (Fig. 1). Leaf growth is one of the most drought-sensitive processes (Boyer, 1970) and leaf elongation rate (LER) response to water deficit has been observed in several grasses (Bouchabke *et al*., 2006; Clifton-Brown & Jones, 1999; Salah & Tardieu, 1996; Schnyder & Nelson, 1988; Parrish & Wolf, 1983; Volenec & Nelson, 1982). Drought is known to induce stomatal closure, which decreases leaf transpiration (Flexas *et al*., 2004) leading to lower water potential (Durand, 2007) (Fig. 1). Reduction in cell size has been well correlated with the reduction of water potential (Bunce, 1977), leading to shorter leaves. In tall fescue, Durand *et al*., (1995) reported a decrease in LER as well as shorter GZ (–60%) and reduced final cell length (–70%) in response to severe drought. Whether the effect of drought on leaf growth is mediated by a decrease in turgor pressure remains unclear. In some studies, the reduction of leaf growth was related to lower turgor pressure in cells of the GZ (Boyer and Potter, 1972; Bouchabke *et al*., 2006), while it has also been reported that turgor pressure remains unchanged during drought (Nonami & Boyer, 1989) or even increased (Onillon, 1993). The stomatal closure induced by drought also reduces photosynthesis (Bradford and Hsiao, 1982; Schulze, 1986; Kaiser, 1987), yet a transient accumulation of water-soluble carbohydrates (WSC) in green tissues and other plant parts may occur as they are not used due to growth cessation (Onillon *et al*., 1995) (Fig. 1). According to many studies, the increase of NSC content may change the osmotic potential, which is a physiological mechanism in response to drought (Bohnert and Jensen, 1996; Hare *et al*., 1998). This osmotic adjustment in response to water deficit can result in maintenance of water uptake to enhance turgor and maintain leaf elongation (Begg & Turner, 1976; Hsiao, 1973), as observed by Wang *et al*., (2008) on tall fescue leaves.

**Fig. 1.**
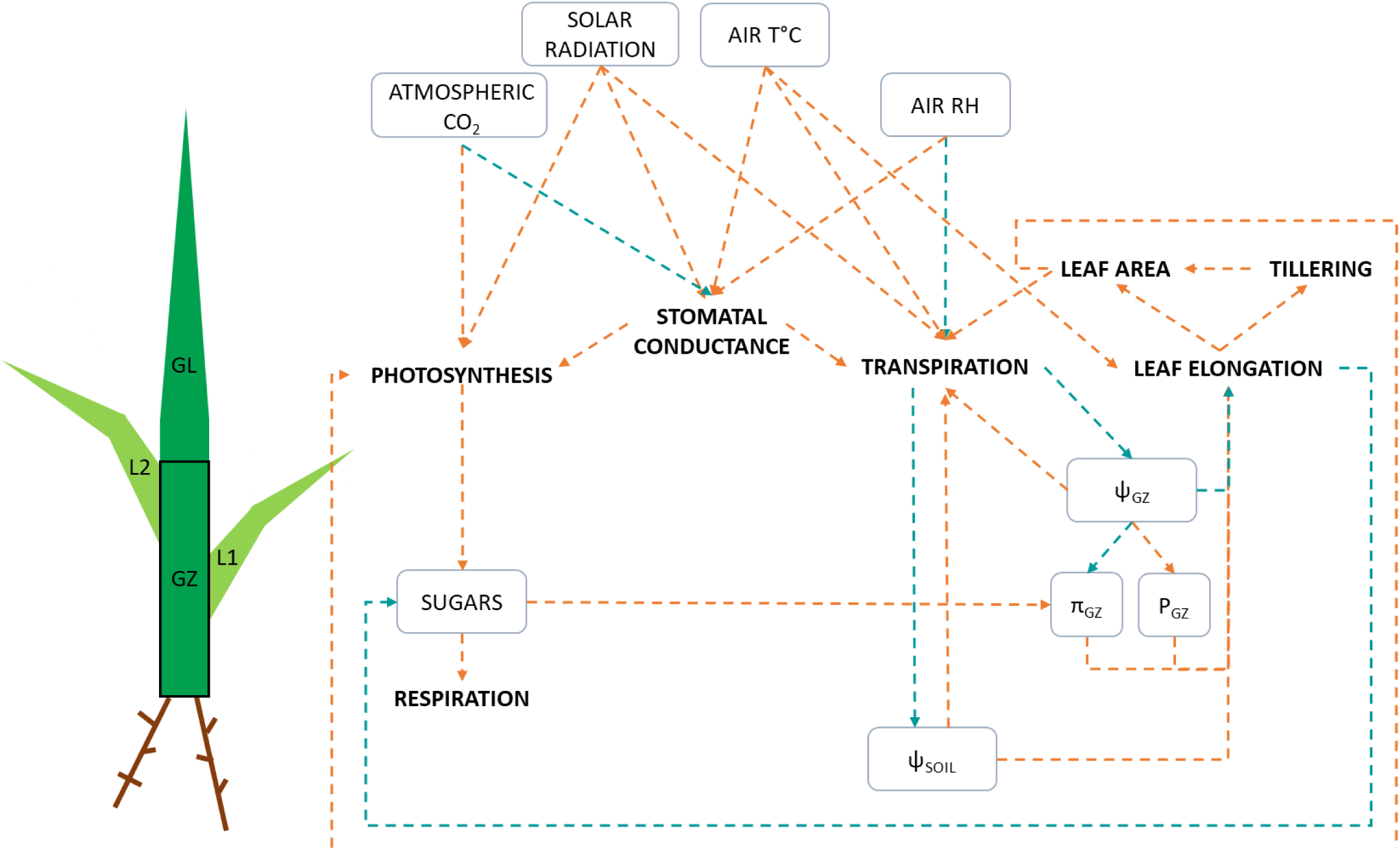
Schematic representation of atmospheric CO_2_ concentration and drought effects on physiological mechanisms of leaf growth and their interrelation in a vegetative grass plant (adapted from Fig. 1 in Baca Cabrera *et al*., 2020). *The plant is represented by the roots, two mature leaves (L_1_ and L_2_), and the growing zone (GZ) of a growing leaf (GL). Orange arrows relates to a positive relationship and blue arrows to a negative one. Leaf elongation is determined by water inflow into the GZ, which is driven by the potential gradient between the GZ and the xylem. The water potential (ψ) is the sum of the turgor potential (P) and the osmotic potential (π). Leaf elongation controls the leaf area and the tillering. Stomatal conductance is sensitive to air relative humidity and temperature, solar radiation and [CO_2_], and represents the physiological control of transpiration. Transpiration is also sensitive to solar radiation, air temperature and relative humidity, as well as the soil water potential. In return, transpiration affects the soil water potential as well as the potential in the GZ. Photosynthesis is directly influenced by [CO_2_], solar radiation and indirectly by the CO_2_ effect on stomatal conductance, and may act on the contribution of water-soluble carbohydrates to the osmotic potential of expanding cells. All of these processes will finally determine the total leaf area which will in turn affect photosynthesis and transpiration*.

It is well known that elevated [CO_2_] stimulates photosynthesis (Gamage *et al.,* 2018; Makino & Mae, 1999) and water-soluble carbohydrate (WSC) production (+46% for wheat at [CO_2_] = 550 ppm according to Thilakarathne *et al*., 2015) (Fig. 1). It has also been shown that increased [CO_2_] led to a reduction of stomatal conductance which could reach ∼40% at [CO_2_] = 700 ppm in herbaceous species (Morison and Gifford, 1984; Field *et al*., 1995; Morison, 1998). This stomata closure can decrease leaf transpiration by 50% (Poorter *et al*., 2022), preserving soil water for a longer period (Fig. 1). This results in an increase of cell division and expansion (Ranasinghe and Taylor, 1996) and of LER by ∼35% for wheat grown at [CO_2_] = 700 and 550 ppm, compared with ambient [CO_2_] (Seneweera & Conroy, 2005; Thilakarathne *et al*., 2015 respectively). Studies also reported an increase in shoot dry mass by 20% in C3 species (Ainsworth and Long, 2005) and of leaf area by ∼40% on wheat at [CO_2_] = 700 ppm compared with ambient [CO_2_] (Morison & Gifford, 1984; Thilakarathne *et al*., 2015) (Fig. 1).

The interactive effect of [CO_2_] and drought on leaf growth is difficult to predict, as on the one hand, elevated [CO_2_] reduces transpiration per unit leaf area, thus mitigating the effects of drought (Yu *et al*., 2012), and on the other hand, elevated [CO_2_] increases total leaf area and therefore water use. With lower stomatal conductance and transpiration rate, water is withheld and soil dries more slowly at elevated [CO_2_], as shown on wheat grown at [CO_2_] = 700 ppm (Bunce, 1998). Under drought conditions, higher water potential is maintained and osmotic potential declines more rapidly at elevated rather than ambient [CO_2_] level (Tyree and Alexander, 1993). It has been observed that the less negative water potential at elevated [CO_2_] allowed plants to wilt less and to incur less drought related damage than plants did in the ambient [CO_2_] (on wheat grown at [CO_2_] = 1000 ppm by Sionit *et al*., 1981; on perennial ryegrass by Nijs *et al.,* 1989; for several C3 and C4 grasses by Sionit & Patterson, 1985). With higher photosynthesis rate at elevated [CO_2_], plants may accumulate more WSC throughout stress periods and therefore maintain higher turgor pressure (Sionit *et al*., 1980). The increased level of WSC enhances development of new tissues during a drought (Wall *et al*., 2006). However, mitigation effects of water stress by elevated [CO_2_] are not always observed. In perennial grassland grown under [CO_2_] = 560 ppm, plant biomass decreased by 37% when both rainfall and nitrogen were at their lowest level (Reich *et al*., 2014). In the study of Baca Cabrera *et al*. (2020), ryegrass plants grown at [CO_2_] = 800 ppm and low VPD = 0.59 kPa, transpired less (–55%) than at ambient [CO_2_], but there was no effect of CO_2_ x VDP interaction neither on water, osmotic and turgor potentials, nor WSC in mature leaves. Elevated [CO_2_] and water deficit affect leaf growth of most C3 species; however there is considerable variation in their magnitude across species highlighting genetic variability and diversity in the experimental design and measurement methods (Kimball and Idso, 1983; Poorter, 1993; Idso, 1994; Kimball, 2016). With the contrasting effects reported in the studies, a global view of the leaf growth response does not yet exist.

To explore these unknowns, we conducted an experiment in growth chambers, with two different [CO_2_]: 200 ppm (‘half ambient’, close to the [CO_2_] in which Poaceae have evolved) and 800 ppm (‘double ambient’, as projected for the end of the century, under a pessimistic scenario of greenhouse gas emission (IPCC, 2023), and also experienced by Poaceae during their early evolution). The two CO_2_ levels were chosen to generate contrasted states in terms of photosynthesis and transpiration in order to assess the potential plasticity of plant morphogeneis. These CO_2_ treatments were combined with watered and drought conditions, in order to evaluate the [CO_2_] x water interaction on transpiration and leaf growth in winter wheat (*Triticum aestivum L.*), an annual cereal; tall fescue (*Festuca arundinacea*) and perennial ryegrass (*Lolium perenne L.*), two major perennial grasses. These species were chosen in order to have contrasting dynamics of morphogenesis (leaf growth rate, tillering ability) and drought tolerance mechanisms related to stomata regulation (Thomas, 1986; Carrow, 1996; Jiang and Huang, 2001; Kemp *et al*., 2001; Cross *et al*., 2013). Cultivars of each species were selected among the ones used in temperate oceanic climate. More specifically, we hypothesised: (1) Despite the negative effects of CO_2_ on stomatal conductance and instantaneous leaf transpiration, total water consumption by the whole plant can be higher at elevated [CO_2_], due to the increase in leaf area. (2) [CO_2_] mitigates the negative impacts of drought on LER and final leaf length. (3) Elevated [CO_2_] reduces leaf transpiration and lowers osmotic water potential through enhanced production of WSC, which mitigates the decrease of nighttime water and turgor potentials in leaf growing zones under drought conditions. (4) Leaf growth responses to [CO_2_] x water stress interaction present genetic variability variations amongst Poaceae species.

## 2. MATERIAL AND METHODS

### 2.1 Experimental design, treatments and growth conditions

For the present study, we grew plants of winter wheat (*Triticum aestivum L*.) *cv.* Soissons, perennial ryegrass (*Lolium perenne L.*) *cv.* Bronsyn and tall fescue (*Festuca arundinacea*) *cv.* Iliade under two levels of CO_2_ x two water supply regimes. The experiment was performed in two growth chambers (97132/7NU, Froids et Mesures, Beaucouzé, France), either set at 200 or 800 ppm of [CO_2_]. In both chambers, CO_2_ was supplied from a tank (CO_2_ 4.5 B50, Linde gas) and mixed with dry CO_2_-free air obtained from a screw compressor (ASD 47 T SFC, Kaeser, Coburg, Germany) and a self-regenerating adsorption dryer (SRE BC 222 GS SP, Chaumeca Gohin, Haubourdin, France). The mixed air was then continuously injected into the chambers by means of mass flow controllers (D-6311 DM and D-6371 DM, Bronkhorst, Ruurlo, Netherlands). CO_2_ concentration in each chamber was measured every 10 sec by an infrared gas analyser (Li-840, Li-Cor, Lincoln, NE, USA), the records are presented in the Supplementary Data (Fig. S3A). The growth chambers were set at 16-h photoperiod with 430 µmol PAR m^-2^ s^-1^ at plant height as provided by HQI lamps (POWERSTAR HQI-T 400D lamps, OSRAM, Munich, Germany). Relative humidity was maintained at 70% and day/night temperatures were set to 19/22°C (Supplementary Data Fig. S3B, C). The temperature was higher during nighttime in order to keep the leaf temperature constant (Supplementary Data Fig. S3D).Plants were grown in single 0.7 L plastic pots (18 x 7 x 7 cm) filled with vermiculite and with 180 g coarse gravel at the bottom. Aluminium foil was placed on the surface of each pot around the plant stem to prevent water loss by evaporation. A total of 480 pots per chamber, including 48 per treatment, were grouped together in containers at a density of 240 plants m^-2^. Each treatment was arranged in two blocks which were surrounded by border plants (Supplementary Data Fig. S2). From seed germination to the emergence of leaf 6 to 7 (42 ± 8 days after sowing according to species and CO_2_ treatments), all plants were irrigated daily with a standard nutritive solution composed of: 1.9 mM KNO_3_, 0.6 mM Ca(NO_3_)_2_, 2.5 mM NH_4_NO_3_, 0.2 mM CaCl_2_, 0.1 mM NaCl, 0.1 mM MgSO_4_, 0.4 mM KH_2_PO_4_, 0.1 mM K_2_HPO_4_, 50 µM H_3_BO_3_, 1 µM CuSO_4_, 4 µM MuSO_4_, 4 µM ZnSO_4_, 1 µM Na2MoO_4_, 100 µM FeSO_4_, 110 µM HEDTA. The quantity of solution provided was regulated so in order to maintain a Soil Relative Water Content (SRWC) between 0.75 and 1. Firstly, irrigation was programmed identically between the two [CO_2_] treatments, and then adapted according to plant development. From the emergence of leaf 6-7, half of the plants was irrigated as before (watered treatment) while the water supply was withheld for the other half (unwatered treatment). The experiment stopped when leaf elongation rate (LER) of unwatered plants was null (53 ± 7 days after sowing according to species and CO_2_ treatments) (Supplementary Data Table S1).

### 2.2 Plant transpiration and soil relative water content

Individual and instantaneous plant transpiration rate (g H_2_O min^-1^ plant^-1^) was calculated every minute gravimetrically, by using a system of load cells placed under 10 plants chosen randomly among the 48 plants per CO_2_ x water x species treatment (Supplementary Data Fig. S1).

The soil relative water content (SRWC) was calculated for the plants placed on the load cells. For each plant *i*, SRWC was calculated for each day *j*, as the cumulative difference between water supply from *j* = 0 (W_irrigation_, g H_2_O plant^-1^) and water loss through transpiration from *j* = 0 (E, g H_2_O plant^-1^), divided by the field capacity (g H_2_O):

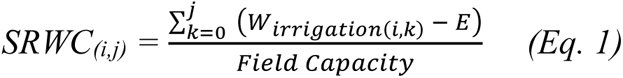

Field capacity was calculated for ten pots filled with vermiculite and coarse gravel as in the growth chambers and was 189 ± 6 g.

### 2.3 Leaf elongation rate (LER)

All plants were in the vegetative stage and depending on the species, were bearing from 6 to 11 leaves on the main stem at the end of the experiment. Leaf length was measured daily with a 1-mm accurate ruler on the main stem of the plants placed on the load cells (10 per treatment). Measurements were performed at the end of the light period on every visible growing leaf (one or two leaves per tiller), by recording the distance between the tip of the elongating leaf and the ligule of the youngest fully expanded leaf. Below, LER are presented at tiller level (cm tiller^-1^ day^-1^) by cumulating the elongation of all visible leaves of the main stem (Gastal *et al*., 1992).

### 2.4 Haun index (HI)

The HI (Haun, 1973) was calculated for main stems in order to analyse the differences in plant development between treatments. HI was calculated daily as the number of ligulated leaves (NL) plus the ratio between the length of the emerged growing leaf (LGL) and its final length (LLL):

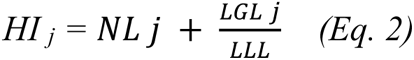

where *j* is the day.

### 2.5 Leaf area and dry matter

Plant sampling was distributed throughout the experiment on two different days: (1) on the first day of the drought treatment, and (2) at growth cessation in the unwatered treatments (LER of plants was null). For two species, ryegrass and tall fescue, we performed a complementary sampling at mid-drought, defined as the time when the LER of unwatered plants was reduced by 50% compared with that of the watered treatment.

Five plants per treatment were sampled on each sampling date, in order to measure plant dry matter and its allocation between leaves, stems and roots. On the three sampling days, the blades of the youngest three leaves on the main stems were collected and scanned using the IMAGE J software (Schneider *et al*., 2012). Leaf area and Specific Leaf Area (SLA, cm² g^-1^) were then calculated on the three sampling days from these samplings. The biomass of roots, blades and sheaths of the main stem, as well as those of the other tillers of the plant, were dried at 60°C for three days. Samples were weighed on a precision balance (BP121S, Sartorius, Sartorius AG 37 075, Göttingen, Germany) and grounded in a vibratory mill (MM200, Retsch, Retsch-Allee 1-5, Haan, Germany).

### 2.6 Leaf water potential and osmotic potential

On each of the three sampling days described above, osmotic and water potentials of the growing zone (GZ) were measured on five plants per treatment, in the middle of nighttime periods. Water potential was measured on the first 5 mm located at the basis of the GZ of the main stem by means of psychrometer chambers (C-52, Wescor Inc., Logan, UT, USA). The next five 15 mm long of the GZ samples were grouped together by treatment, then placed in a tube, frozen in liquid nitrogen and stored at –20°C. Samples were then brought to ambient temperature for 30 mn and the sap was expressed after thawing under pressure in a syringe. Mean osmotic potential was measured using a vapour pressure osmometer (5600, VAPRO, Wescor Inc., Logan, UT, USA).

### 2.7 Water-soluble carbohydrates (WSC)

WSC concentration was measured on blade, sheath and roots of each sampled plant. 60 mg ground dry material were weighed into 15 mL capped tubes containing a mixture of Brij and 5 mL demineralised water heated to 40°C. Tubes were sealed immediately and transferred to a heat chamber at 40°C for one hour. Then, the tubes were centrifuged at 16000 rpm for 2 min and 2.5 mL supernatants were sampled with a pipette. WSC were analysed with a continuous flow system similar to the one described by Wolf & Ellmore (1975). An aliquot of the WSC extract was hydrolysed in 0.1M sulfuric acid (0.2N) for 40 min at 100°C. After cooling, the reducing power of the hydrolysed carbohydrates was assayed by an Auto-Analyzer (model AAII-02, Technicon Instruments Corp., Tarrytown, USA).

### 2.8 Stomatal conductance to water (gs_w_)

At the end of the experiment, both stomatal conductance in ambient CO_2_ of each chamber and stomatal conductance response to CO_2_ were measured with a Li-6800 (Li-Cor, Lincoln, Nebraska, USA) portable CO_2_/H_2_O gas exchange system with a clamp-on leaf cuvette on five plants per [CO_2_] treatment under well-watered conditions. Leaves of unwatered plants were too small and dried to be measured. Stomatal conductance to water vapour (gs_w_, in mol H_2_O m^-2^ s^-1^) was measured one hour after switching on the lamps using the A-Ci program of the Li-6800 device. The midsection of the youngest fully developed leaf blades of one tiller was enclosed for 30 minutes in the 2 cm² leaf cuvette set at PAR=1500 μmol m^-2^ s^-1^, leaf temperature = 20°C and VPD = 1 kPa. gs_w_ response to CO_2_ was measured from 0 to 1 800 ppm in 10-12 points.

### 2.9 Statistics

The CO_2_–induced change of each studied variables *X* was calculated as: 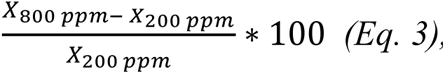, and in the same way, the drought-induced change of *X* by: 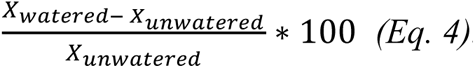. Exploratory data analysis, analysis of variance and linear regression techniques were performed with R v.4.0.2 (R Core Team, 2019), and figures were created using the ggplot2 package (Wickham, 2016). Analysis of variance (ANOVAs) was performed following a two-factor linear model such as follows:

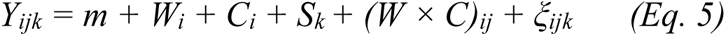

where *Y* is any dependent variable, *m* the mean value of *Y*, *W_i_* the effect of water treatment *i*, *C_j_* the effect of [CO_2_] treatment *j, S_k_* the effect block *k,* and *ξ* the random error of measurement *ij*.

Normal distributions of the residuals of ANOVAs were tested using the Shapiro–Wilk test and homoscedasticity was checked using the Levene test. T-test was used for multiple pairwise comparisons between groups. Means of different treatments were tested with the least significant difference at a probability of 0.05.

The linear correlation coefficient (Pearson) was used to calculate the dependence between two quantitative variables.

## 3. RESULTS

### 3.1 Effects of water deficit and [CO_2_] interaction on water use

Plant cumulated transpiration (Fig. 2A) increased with plant development and reached 350 to 800 g per plant at the end of the experiment. Overall, the total transpiration of tall fescue plants (∼600 g) was greater than that of ryegrass (∼490 g) and wheat (∼500 g). After drought onset, cumulated transpiration of plants rapidly slowed down (in ∼1.5 days) and at the end of the experiment, the total amount of water transpired was ∼70% lower compared with watered conditions (p-value<0.5). [CO_2_] showed contrasted effects on cumulated transpiration according to the species. For tall fescue and wheat, if any, [CO_2_] had a slight effect, as total plant transpiration of watered plants was on average 7% lower at 800 ppm than at 200 ppm (not statistically significant). Conversely, elevated [CO_2_] drastically increased cumulated plant transpiration of ryegrass, which was 40% and 20% higher at 800 ppm than at 200 ppm for watered and unwatered plants, respectively (p-value<0.001 and <0.05).

**Fig. 2.**
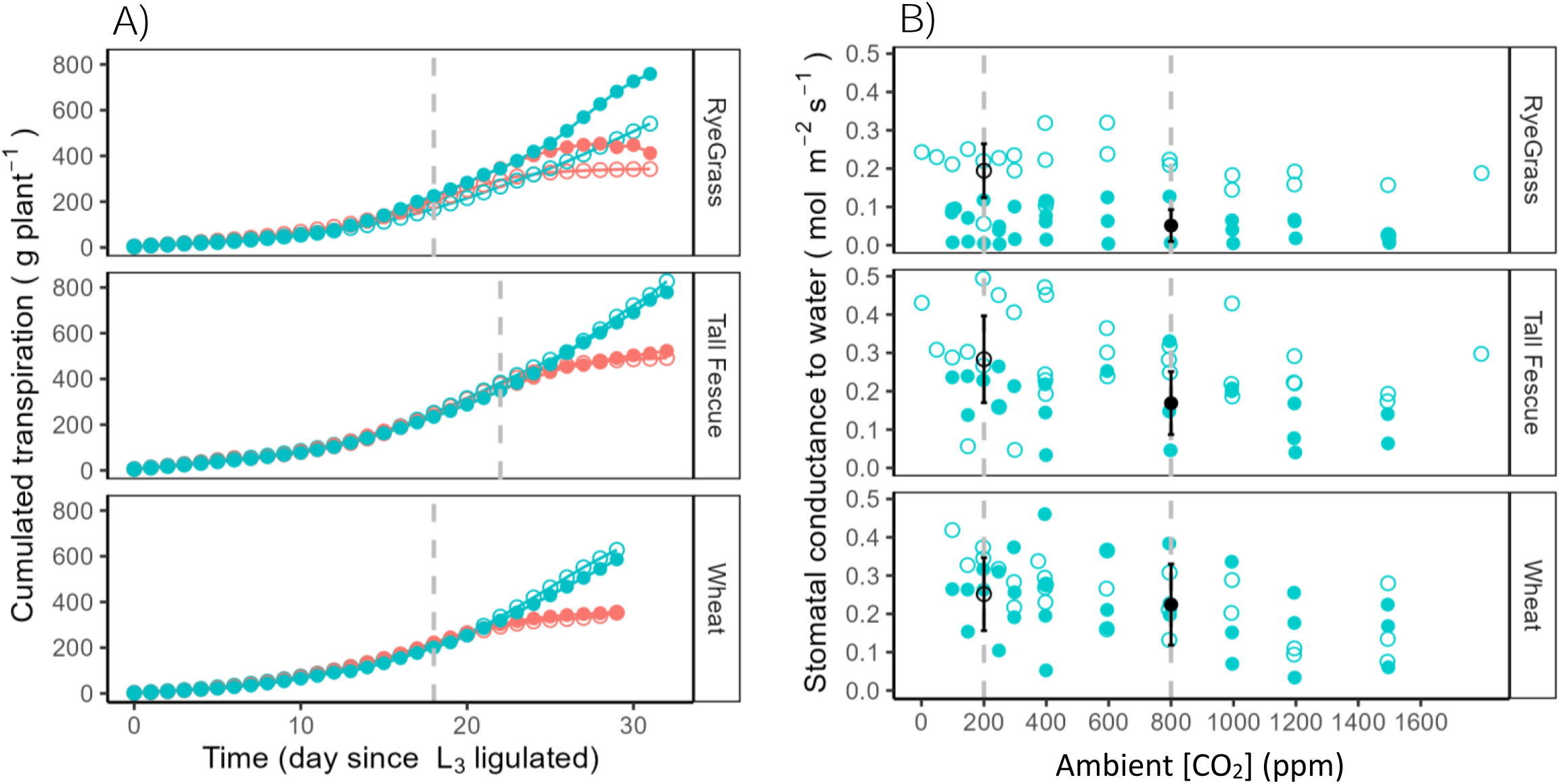
Daily cumulated transpiration (g) per plant (A), and stomatal conductance to water (mol m^-2^ s^-1^) as function of ambient [CO_2_] (ppm) (B) for *Lolium perenne L.* (upper panels), *Festuca arundinacea* (mid-panels) and *Triticum aestivum L.* (bottom panels). *Results are shown for plants grown at 200 (open circles) or 800 ppm [CO_2_] (closed circles) in watered (blue) or unwatered (red) conditions. Plant transpiration was calculated gravimetrically and averaged on ten plants per treatment. The beginning of the drought treatment is shown by the vertical dotted line. Stomatal conductance (gs_w_) response to ambient [CO_2_] (Ca) was measured for watered plants only (n = 5). Black circles and their associated error bars are mean ± standard deviation data of gs_w_ for ambient [CO_2_] (Ca) at 200 and 800 ppm*.

Stomatal conductance to water (gs_w_) in each chamber as well as gs_w_ response to CO_2_ were measured at the end of the experiment on leaves of watered plants (Fig. 2B). Overall, gs_w_ was lower for ryegrass than for tall fescue and wheat (p-value<0.05). In contrast with the observations for whole plant transpiration, plants grown at elevated [CO_2_] showed significantly lower gs_w_ than those grown at 200 ppm, for ryegrass and tall fescue (–70% and –30% respectively, p-value<0.05). For wheat, the gs_w_ did not respond significantly to ambient [CO_2_] (p-value>0.05). There was no significant adaptation of the sensitivity of the gs_w_ to CO_2_ in wheat during this experiment.

### 3.2 Effects of water deficit and [CO_2_] interaction on plant growth

The responses of cumulated plant transpiration to [CO_2_] were strongly related to leaf area development (Fig. 3A). Leaf area of watered plants increased with time, reaching 130 cm² to 360 cm² at the end of the experiment. Drought reduced leaf area by ∼170% at the end of the experiment, all treatments taken together (p-value<0.05). Elevated [CO_2_] increased leaf area of watered plants by 40% in wheat, by 70% in tall fescue and by 180% in ryegrass. Under drought conditions, leaf area of unwatered plants increased until mid-drought at [CO_2_] = 800 ppm, and then decreased due to senescence (by 40% in tall fescue and 170% in ryegrass); whereas at [CO_2_] = 200 ppm, drought immediately affected leaf area which remained almost constant at the different growth stages.

**Fig. 3.**
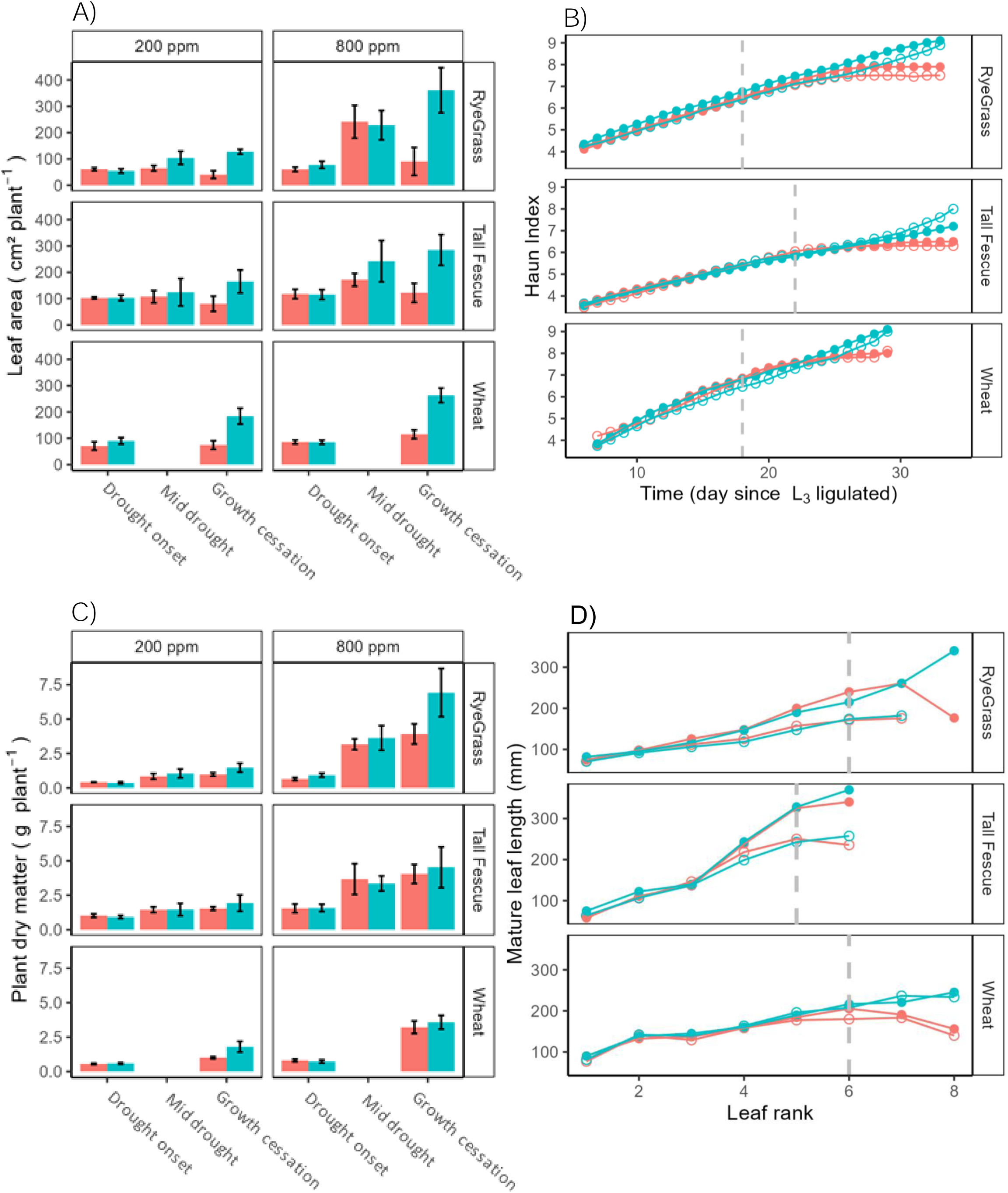
Leaf area (cm²) per plant (A), Haun index (B), Plant dry matter (g) per plant (C), and and final length of ligulated leaves of the main stem (mm) as function of the leaf rank (D) for *Lolium perenne L.* (upper panels), *Festuca arundinacea* (mid-panels) and *Triticum aestivum L.* (bottom panels) as function of drought progression. *Results are shown for plants grown at 200 (open circles) or 800 ppm [CO_2_] (closed circles) in watered (blue) or unwatered (red) conditions. The start of the drought treatment is shown by the vertical dotted line. Data is averaged on five plants per treatment. Error bars represent the mean ± standard deviation. Plant dry matter is composed of main stem, tillers and roots dry matter*.

Plant dry matter (PDM) was dramatically affected by water deficit and [CO_2_] (p-value<0.05). PDM increased with time in watered conditions and finally reached 1.5 to 7 g. After drought onset, PDM reached a plateau and was ∼25% lower for unwatered plants than for watered plants (Fig. 3C). Elevated [CO_2_] increased DM of watered plants which was ∼207% higher than that at 200 ppm at the end of the experiment. We observed an increase of the main stem DM by ∼80% while the DM of other tillers (plant DM – main stem DM) increased by ∼280% between 200 and 800 ppm, for watered plants. Therefore, the enhancement of plant DM with [CO_2_] was mainly due to a dramatic increase in other tillers biomass which can be explained by an increase in tiller number and individual tiller DM (Supplementary Data Fig. S5A). [CO_2_] effect persisted under drought conditions, as the decline of PDM was higher at 200 than 800 ppm: ∼-30% at 200 ppm, and ∼-20% at 800 ppm between unwatered and watered plants of the three species.

During the experiment, Haun Index (HI) increased faster in wheat and ryegrass than for tall fescue (Fig. 3B). On average, wheat and ryegrass produced one leaf every ∼5 days, whereas it took ∼7 days for tall fescue (p-value<0.05). After drought onset, HI progressively reached a plateau and did not increase further than ∼7.3, as no more leaves emerged. [CO_2_] did not affect HI for all three species on both water regimes (non-significant p-value).

Final leaf length (Fig. 3D) increased with leaf rank in watered conditions (p-value<0.05). Tall fescue showed the longest mature leaves on L6 stage, higher by ∼+30% all treatments combined. In watered conditions, elevated CO_2_ increased final leaf length by 40 and 30 % for tall fescue and ryegrass respectively (p-value<0.05) for which leaves were ∼40 and 30% longer at 800 than 200 ppm (on L6 stages for tall fescue and ryegrass respectively, p-value<0.05). In contrast, wheat leaves had similar length at both [CO_2_] (non-significant p-value). In unwatered condictions, final length of mature leaves was reduced by ∼10% at both [CO_2_] (p-value<0.05), for all three species. Furthermore, [CO_2_] did not affect plant response to drought (p-value>0.05).

### 3.3 Effects of water deficit and [CO_2_] interaction on leaf growth

With drought onset, Leaf Elongation Rate (LER) of unwatered plants decreased rapidly compared with watered plants, reaching 0 cm day^-1^ after 9 to 12 days for tall fescue, 10 to 13 days for wheat, and 11 to 14 days for ryegrass (Fig. 4A). [CO_2_] had a positive effect on the LER of plants subjected to drought. Unwatered plants grown at elevated [CO_2_] were able to maintain leaf elongation for a longer time than those grown at [CO_2_] = 200 ppm: three more days for ryegrass and tall fescue, and two more days for wheat (p-value<0.05).

**Fig. 4.**
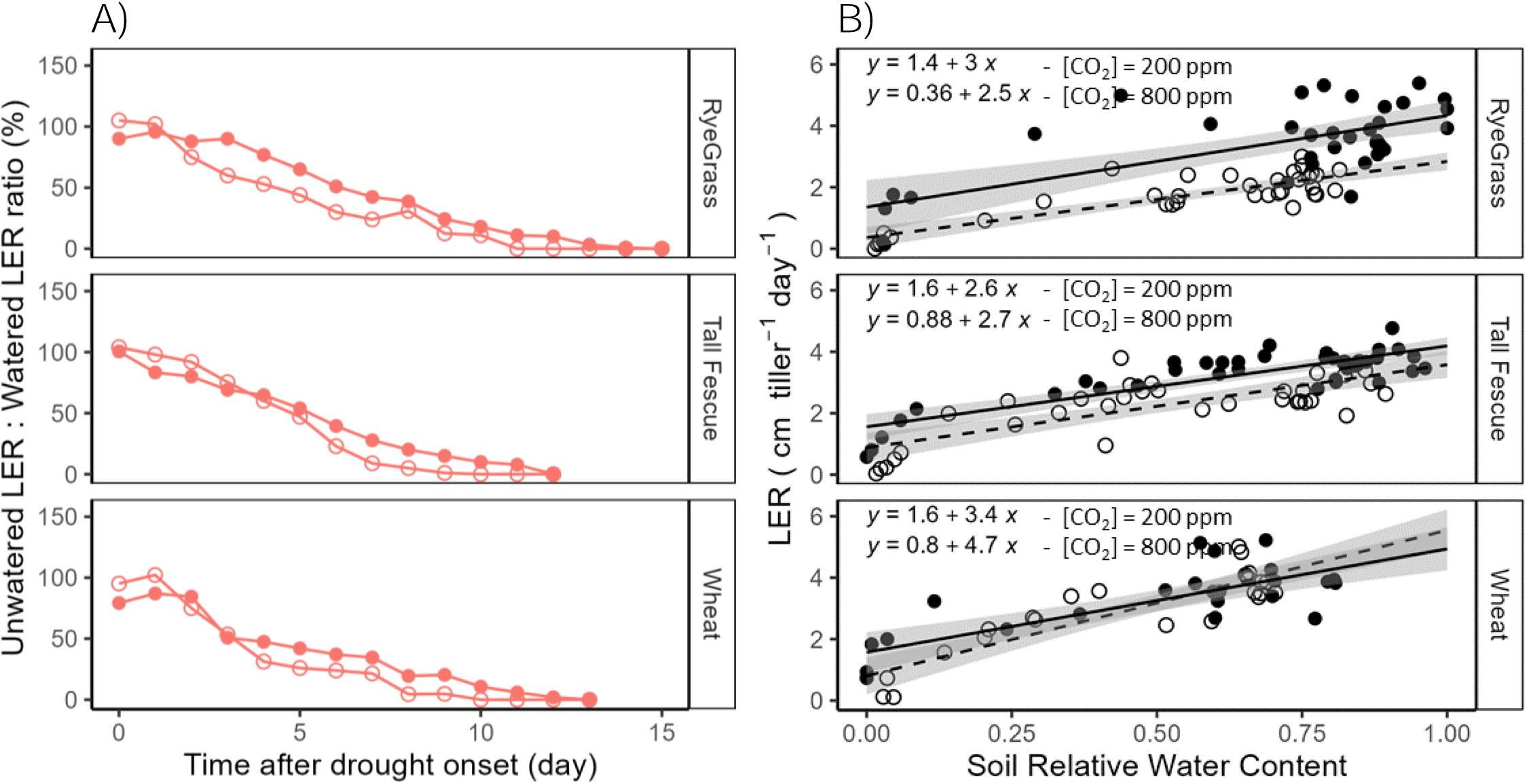
LER Unwatered: LER Watered ratio as function of time after drought onset (day) (A), and LER (cm tiller^-1^ day^-1^) as function of the SRWC (B) for *Lolium perenne L.* (upper panels), *Festuca arundinacea* (mid-panels) and *Triticum aestivum L.* (bottom panels). *Results are shown for plants grown at 200 (open circles) or 800 ppm [CO_2_] (closed circles). Lines and equations relate to linear regression with standard error bounds in grey*.

We found a positive linear relation between LER and the Soil Relative Water Content (SRWC) which ranged from optimal water availability (SRWC = 1) to extreme drought (SRWC = 0) (Fig. 4B). For all species, the y-intercept of the linear relation between LER and SRWC was signigicantly different with CO_2_ (p-value<0.05, Supplementary Data Table S2) and was lower at 200 ppm than at 800 ppm. The slope of the equation was not affected by [CO_2_] (p-value>0.05, Table 1). In ryegrass and tall fescue, for a similar level of SRWC, leaves were elongating faster at 800 than at 200 ppm of [CO_2_] (p-value<0.05). In contrast, [CO_2_] showed no statistically significant effect on the relationship between LER and SRWC for wheat. With the highest slope, at both CO_2_ levels, wheat was the species with the LER the most sensitive to SRWC.

### 3.4 Effects of water deficit and [CO_2_] interaction on growing zone water status

WSC concentration ([WSC]) (Fig. 5A; Supplementary Data, Fig. S7A, B) was highest in the sheaths, then in the blades and finally in the roots. The response of WSC being similar to water and [CO_2_] treatments between the different organs, only data of the blades are presented (Fig. 5A). [WSC] of watered plants decreased with time for wheat and tall fescue throughout the experiment, but it tended to increase for ryegrass. At the end of experiment, [WSC] of unwatered plants grown at [CO_2_] = 200 ppm increased by 15% compared to watered plants, while it increased by ∼30% for tall fescue, ∼60% for wheat, and decreased by ∼40% for ryegrass at [CO_2_] = 800 ppm. Elevated CO_2_ resulted in higher [WSC] by ∼60% for watered plants of all three species. With drought, [WSC] was about ∼50% higher at 800 than 200 ppm for ryegrass and tall fescue, and ∼15% higher for wheat at drought onset, and increased with time to finally be ∼80% higher at 800 ppm for wheat and tall fescue, and ∼40% for ryegrass at the end of the experiment (p-value<0.05 for ryegrass and tall fescue).

**Fig. 5.**
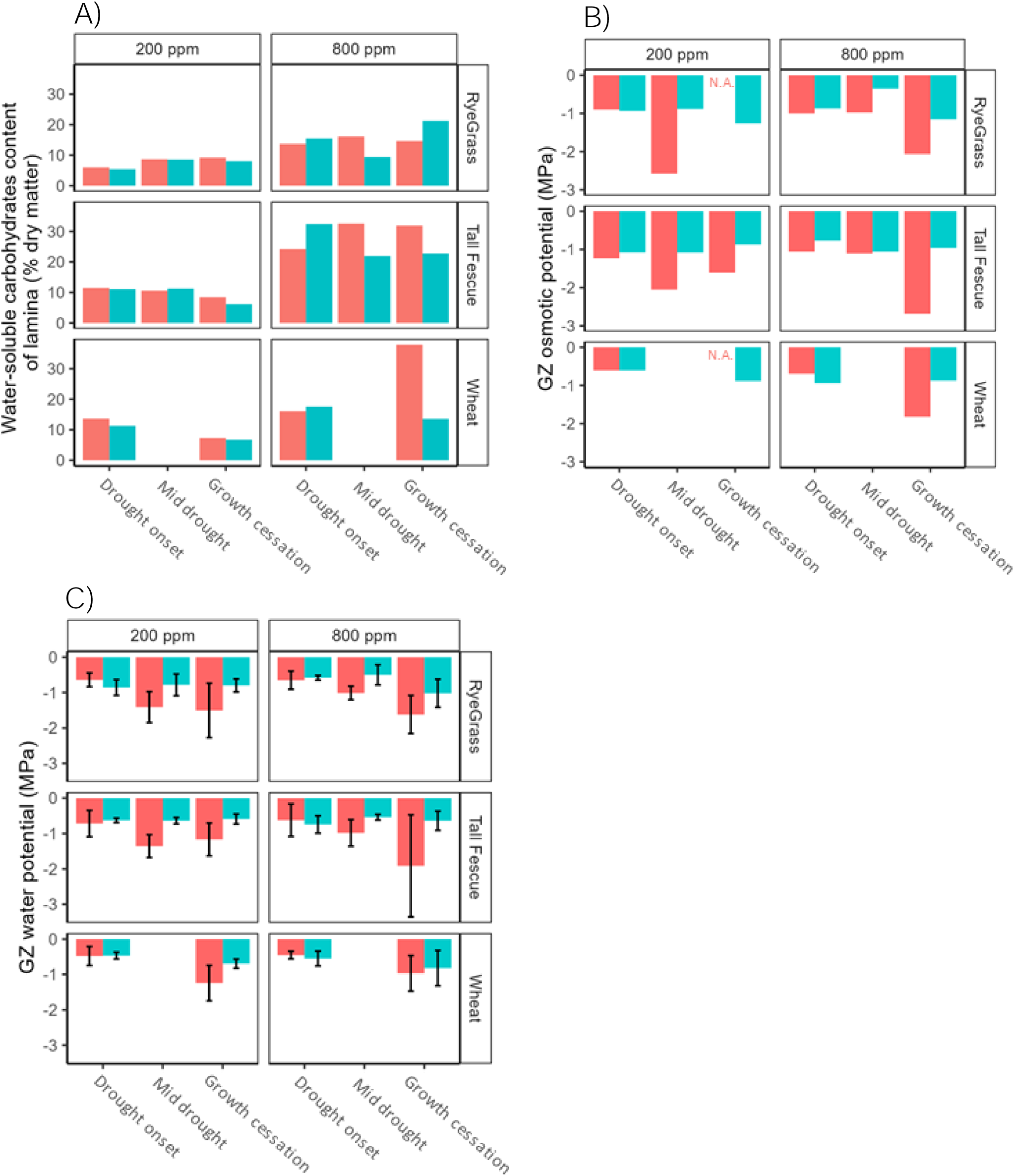
Water-soluble carbohydrate content (% dry matter) per plant (A), Growing zone osmotic potentials (π _GZ_ MPa) (B), and Growing zone water potentials (ψ _GZ_, MPa) (C) for *Lolium perenne L.* (upper panels), *Festuca arundinacea* (mid-panels) and *Triticum aestivum L.* (bottom panels) at three drought stages: drought onset, mid-drought and growth cessation. *Results are shown for plants grown at 200 (left side panel) or 800 ppm [CO_2_] (right side panel) in watered (blue) or unwatered (red) conditions. Data is averaged on five plants per treatment*.

Osmotic potentials of growing zone (π_GZ_) of watered plants remained relatively constant: ∼-0.9 MPa at [CO_2_] = 200 ppm for all three species and ∼-1 MPa at 800 ppm for ryegrass and tall fescue, and ∼-0.7 for wheat (Fig. 5B) and was not significantly different according to species. We observed more variability under unwatered than watered conditions between species and [CO_2_]. Nevertheless, π_GZ_ of unwatered ryegrass and wheat could not be measured at 200 ppm at the end of the experiment, as the plants were too small, which made the extraction of GZ difficult. After drought onset, for all species π_GZ_ decreased from mid-drought (except for tall-fescue at [CO_2_] = 800 ppm) and was ∼50% lower at the end of the experiment at both [CO_2_] compared with watered conditions. However, [CO_2_] did not significantly affect π_GZ_ during nighttime in all species (non-significant p-value). The osmotic potential and the water-soluble carbohydrate content were not correlated (corr<0.5, Supplementary Data Table S3).

For watered plants, water potential of growing zone (ψ_GZ_) ranged from –0.8 to –0.6 MPa throughout the experiment (Fig. 5C) and showed no significant difference according to species. Under drought, ψ_GZ_ decreased for all three species (p-value<0.05) and was lower compared with watered conditions from mid-drought time for ryegrass and tall fescue (p-value<0.05). Results showed more disparity between species and [CO_2_] under unwatered than watered conditions. [CO_2_] did not affect ψ_GZ_ of watered plants during nighttime (non-significant p-value). Under drought conditions, ψ_GZ_ did not respond to [CO_2_] either (non-significant p-value), however we observed interspecific variations. At the end of the experiment, while ψ_GZ_ of wheat dropped by ∼40% and ∼15% (non-significant p-value), ψ_GZ_ of tall fescue dropped by ∼50% and ∼70% (non-significant p-value), and ψ_GZ_ of ryegrass dropped by ∼50% and ∼40% (non-significant p-value), at [CO_2_] = 200 and 800 ppm respectively. [CO_2_] slowed down the ψ_GZ_ decrease as ψ_GZ_ were higher at [CO_2_] = 800 ppm than at 200 ppm at mid-drought time.

Variation of nighttime ψ_GZ_ was highly dependent on SRWC with similar pattern for the three species, i.e., ψ_GZ_ remained stable in wide range of SRWC and dropped sharply when the soil dried (Fig. 6). Two linear regression models were used to estimate the threshold from which ψ_GZ_ dropped. The response of ψ_GZ_ to SRWC was significantly different between [CO_2_] treatment for ryegrass (Supplementary Data Table S4) while it was not affected for tall fescue and wheat (non-significant p-value). At [CO_2_] = 200 ppm, ψ_GZ_ of ryegrass started to fall from SRWC = 0.26, while the relation was more linear at [CO_2_] = 800 ppm. For tall fescue and wheat, a threshold could be defined at SRWC = 0.19 and 0.27 for tall fescue and wheat respectively, at both CO_2_ levels. Therefore, ψ_GZ_ of wheat was the most responsive to SRWC, followed by ryegrass grown at low CO_2_, tall fescue and finally ryegrass at elevated CO_2_.

**Fig. 6.**
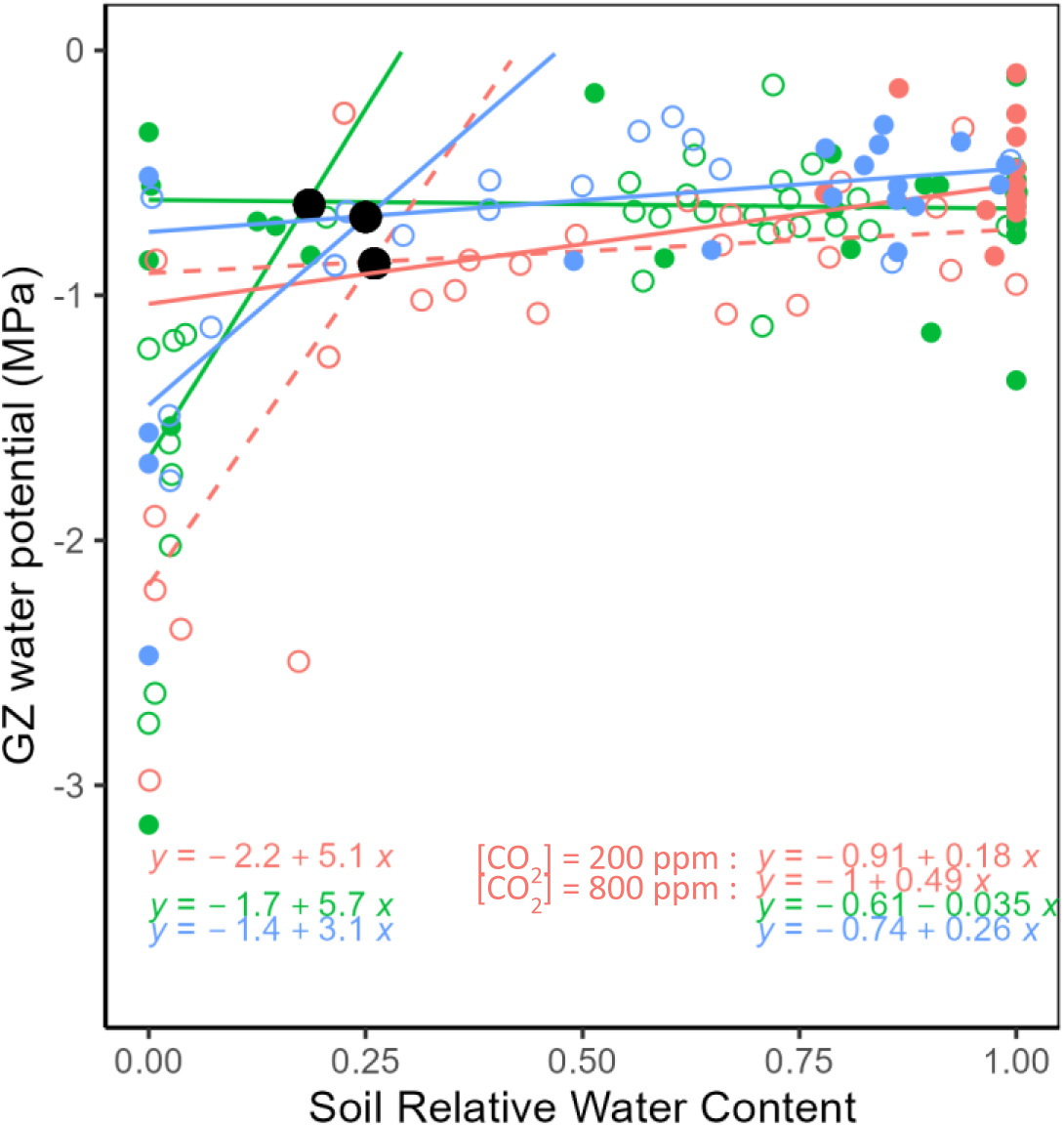
Growing zone water potential (ψ _GZ_, Mpa) as function of Soil Relative Water Content for *Lolium perenne L.* (red circles), *Festuca arundinacea* (green circles) and *Triticum aestivum L.* (blue circles). *Results are shown for plants grown at 200 (open circles) or 800 ppm [CO_2_] (closed circles). The dark circles represented the threshold from which ψ_GZ_ decreased as the soil is drying. Lines and equations relate to linear regression of the ψ_GZ_ treshold for lowest SRWC range (left side) and highest SRWC range (right side) of the ψ _GZ_ threshold*.

## 4. DISCUSSION

### 4.1 Increasing [CO**_2_**] from 200 to 800 ppm did not save water but strongly increased leaf area

We investigated the ability of [CO_2_] to prevent water loss by plants. Consistent with this hypothesis, gs_w_ was lower at elevated [CO_2_] although the intensity of the stomatal closure was markedly different between species (–70% for ryegrass, –30% for tall fescue and –10% for wheat compared with low [CO_2_]). Similar results were observed on ryegrass by Baca Cabrera *et al*. (2020) who reported a decrease in gs_w_ by 75% between [CO_2_] = 200 and 800 ppm, and on tall fescue where gs_w_ decreased by 25% between [CO_2_] = 400 and 800 ppm (Yu *et al*., 2012). For wheat, Bunce (2000) observed a stronger reduction of gs_w_ (50%) between 350 and 700 ppm in open-topped chambers than the one we measured in the present study between 200 and 800 ppm in growth chambers, presumably related to lower temperatures in their field conditions. gs_w_ was not measured in drought conditions in our experiment, nonetheless, it has been reported by (Li *et al*., 2004) that elevated CO_2_ reduced gs_w_ more in droughted treatments (−51%) than in well-watered treatments (−41%) for wheat plants grown at [CO_2_] = 350 and 700 ppm. During the early stages of development, the stomatal closure induced by elevated [CO_2_] leads to a decrease in instantaneous transpiration rates (Supplementary Data Fig. S4A) in our experiment, which was also reported in tall fescue (Chen *et al*., 2015: between 400 and 800 ppm) and wheat (Tyree and Alexander, 1993: between 350 and 800 ppm).

Concominantly, elevated [CO_2_] increased leaf photosynthesis and WSC content in the different organs of the three species. Similar results have been observed on tall fescue by Chen *et al*. (2015), wheat (Thilakarathne *et al*., 2015), as well as other C3 species (Ainsworth and Rogers, 2007). As a consequence, WUE (Supplementary Data Fig. S4B) was higher by ∼60% at elevated CO_2_ for all three species (p-value<0.05). It resulted that the positive effect of [CO_2_] on both WSC accumulation and water conservation enhanced the growth potential, which increased leaf area production with strong interspecific variability. Total cumulated transpiration was either similar, for wheat and tall fescue, or higher for ryegrass at elevated CO_2_ at the end of the experiment.

### 4.2 Elevated [CO**_2_**] mitigated drought effect on leaf growth

It has been shown that transpiration largely affects soil water avalibility but also leaf elongation rate, e.g. during day/night variations (Parrish and Wolf, 1983) or in the case of induced stomatal opening by blue light irradiance (Barillot *et al*., 2020). The decrease of transpiration rate leads to an increase in water potential gradients within the tiller, which induces an increase in the water flow in the growing zone, and therefore a faster elongation. We can therefore suppose that CO_2_-induced stomatal closure lead to an increase in LER. Accordingly, we found that the response of LER to soil water availibity was positively affected by [CO_2_] in ryegrass and tall fescue (Fig. 4B), the two species that showed a large reduction of gs_w_ in response to elevated [CO_2_]. LER of wheat probably did not respond to [CO_2_], due to its lower response of gs_w_. Seneweera & Conroy (2005) observed an accelerated LER by 32% on wheat plants grown at 700 ppm compared with 370 ppm, under well-watered conditions. These contrasting results may be explained by different methods of LER calculation, which was restricted to the 6^th^ leaf for 12 hours in Seneweera & Conroy (2005), while LER was monitored at tiller scale (see section 2.3) for several weeks in the present study.

Our study showed that over a long development time, [CO_2_] delayed the effects of drought on the LER in all species. With the decrease of the transpiration rate which delayed the depletion of soil water, [CO_2_] lowered the rate of decline of SRWC during drought. Soil water conservation has also been observed by Robredo *et al*. (2007) in barley, where soil drying was delayed by 3-4 days at [CO_2_] = 850 ppm compared with ambient [CO_2_]. Similar results were reported for soybean Rogers et al. (1984), maize (Samarakoon & Gifford, 1995) and rice (Vu et al., 1998). If an elevated [CO_2_] allowed the maintenance of growth for a longer period during water stress, it did not prevent the cessation of growth in severe and long drought conditions.

### 4.3 CO**_2_** did not affect nighttime water potential nor osmotic potential of the GZ

The measurement of water potential made it possible to define water status at plant level and to discuss the relationships between the water variables and growth. As [CO_2_] affected LER response of ryegrass and tall fescue to drought, we investigated whether this resulted in an increase in the water potentials of the leaf GZ (ψ_GZ_). This present work provided the first data on water potential of GZ in these conditions. We did not find any [CO_2_] effect on ψ_GZ_, whatever the species and water supply treatment. In addition, the response of ψ_GZ_ to SRWC was not significantly affected by [CO_2_] for tall fescue and wheat. CO_2_ effect on ψ_GZ_ and SRWC relation in ryegrass was likely linked to the stronger response of gs_w_ of this species. It should be noted that in the present study, water potential measurements were performed during nighttime, meaning that the transpiration rate was very low and growth rate likely higher compared with daytime (i.e on tall fescue: Parrish and Wolf, 1983; Durand *et al*., 1995). Our environmental conditions were close to the ones of Baca Cabrera *et al*. (2020) who reported that the effect of [CO_2_] was only significant during daytime. At present, it does not exist a consensus on how the interaction of CO_2_ and water availability affect the leaf water status during daytime. In fact, opposite effects have even been observed under drought, showing lower midday water potential of mature leaves (ψ_L_) at elevated [CO_2_] on barley (Robredo *et al*., 2007) and wheat (Wall *et al*., 2006), similar ψ_L_ on grasses (Ferris and Taylor, 1995; Baca Cabrera *et al*., 2020), or less negative ψ_L_ values for wheat (Sionit *et al*., 1981).

The present work provides original results of osmotic potential in GZ at two CO_2_ levels and in three different species. As expected, osmotic potential of GZ (π_GZ_) decreased with drought, but [CO_2_] had very little effect on π_GZ_, which is consistent with Baca Cabrera *et al*. (2020) in ryegrass who reported π_GZ_ of ∼-1.5 and – 2 MPa at the end of the night, at both 200 and 800 ppm. In contrast, a decrease in the osmotic potential of mature leaves (π_L_) in response to CO_2_ was observed in unwatered plants of wheat (Sionit *et al*., 1981) and tall fescue (Chen *et al*., 2015) at elevated [CO_2_]. However, the latter measurements were taken from mature leaves, which might be different from that of the GZ. In addition, we found that despite the positive effect of CO_2_ on the WSC of the plant, it did not lead to any osmotic adjustement in the GZ in both watered and drought conditions, suggesting a low contribution of sugars to the osmotic potential. It should be noted that we measured WSC and π_GZ_ in different plant tissues (mature leaves, growing zone, respectively), which could also explain their non-correlation. However, it is very likely that CO_2_ not only increased WSC in blades, sheaths and roots but also in GZ. Similar conclusions were raised by Baca Cabrera *et al*. (2020).

Using our measurments on water and osmotic potentials in, we assessed turgor potential (P) which is the physical force needed to sustain enlargment and therefore has a critical role in cell growth (Hsiao *et al*., 1976). The response of GZ turgor potential (P_GZ_) to CO_2_ and drought showed a large variability between the species, notably at the end of the experiment due to the leaf senescence. With drought, P_GZ_ increased and therefore we cannot relate the decrease of LER through a decrease of P_GZ_. Although there are examples of plants exhibiting a positive response of LER to P in relation to water status (e.g. on wheat Fischer and Sanchez, 1979; on maize Bouchabké *et al*., 2006), there are also examples in which expansion is unrelated to P or is overridden by cell wall properties (Termaat *et al*., 1985). Martre *et al*. (1999) reported that P was systematically lower in expanding cells than in mature cells of tall fescue and P_GZ_ was not correlated with the local growth rate along the growing zone. Without cellular expansion, P_GZ_ does not decrease by relaxation and therefore remained high as in mature leaves. Durand *et al*. (1995) reported that GZ size decreased with drought. As dimensions of samples were kept constant throughout the experiment (5 mm base of the leaf), the tissues were probably not growing anymore at the growth cessation time. Therefore, there was no relaxation, which might explain the higher P_GZ_ measured in the basal part of the leaf under such conditions. Onillon (1993) also reported an increase of P_GZ_ in tall fescue under drought conditions.

### 4.4 Interspecific variability in growth response to [CO**_2_**] and drought interaction

To summarize the plant growth response to CO_2_ and the interspecific variability, we presented the relative variation of each components of morphogenesis studied in this work, between 200 and 800 ppm and in the two water conditions (Fig. 7A, B).

**Fig. 7.**
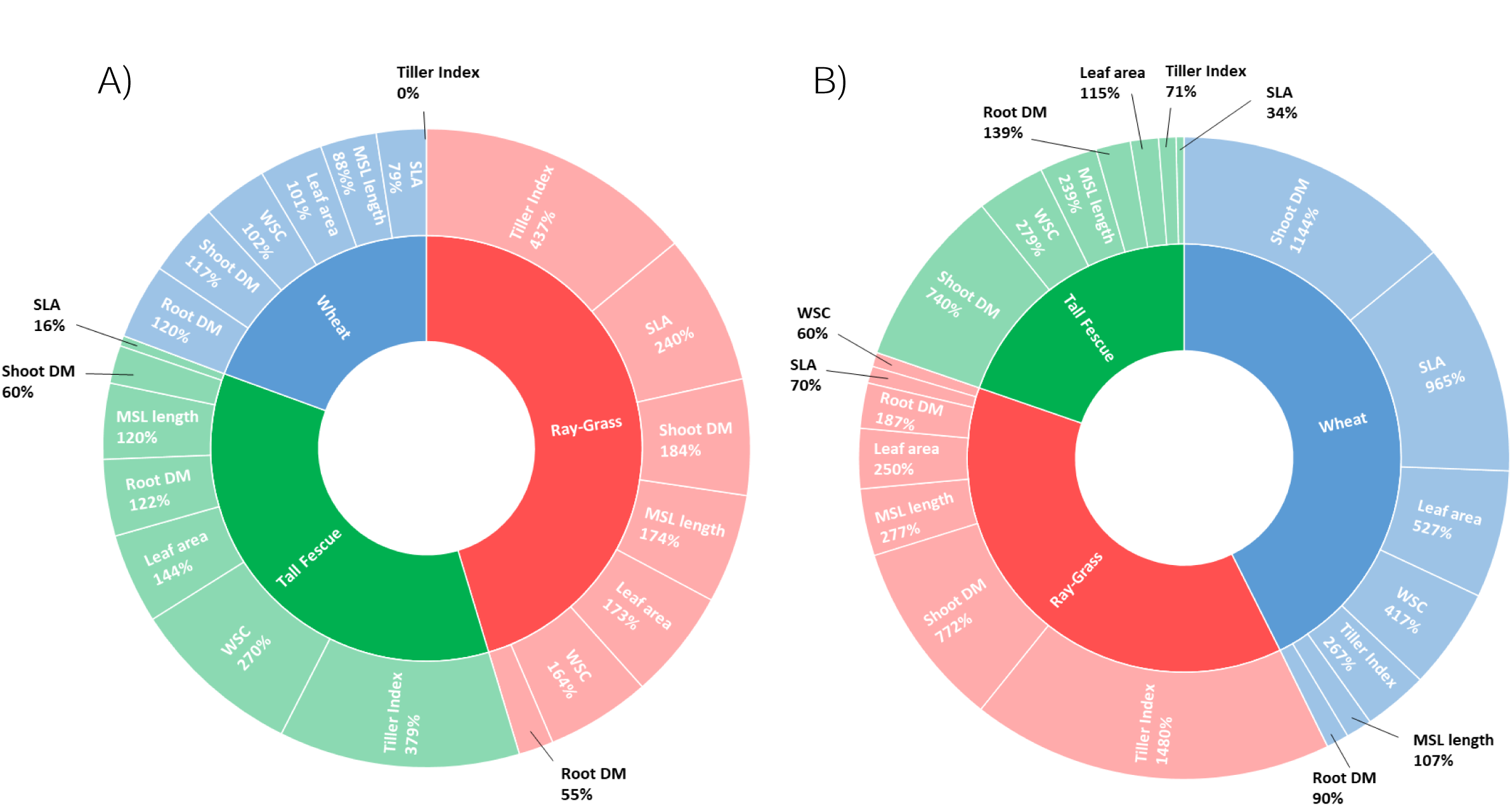
Summary of the CO_2_ effects on plant morphogenesis under well-watered conditions (A), and on plant response to drought (B) for *Lolium perenne L.* (red), *Festuca arundinacea* (green) and *Triticum aestivum L.* (blue). *Results are calculated as the variation between 200 and 800 ppm at the end of the experiment for WSC water-soluble carbohydrates contents (WSC) variable. For the other variables, X, data are the variation between 200 and 800 ppm since drought onset, by:* 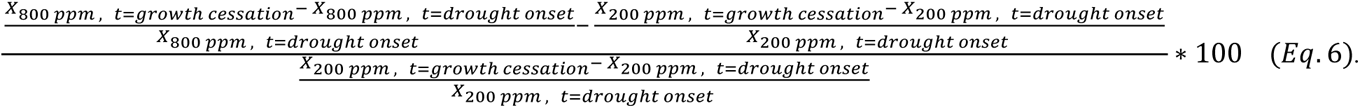 *The size of the variable boxes is proportional to the CO_2_ effect on it. Shoot DM refers to shoot dry matter, root DM to roots dry matter, SLA to specific leaf area, MSL length to total main stem leaves length and Tiller Index is the ratio of dry matter between other tillers and main stem*.

The response of the different components of plant morphogenesis to CO_2_ showed a strong interspecific variability (Fig. 7A). CO_2_ effect was stronger for the perennial forage grasses than for wheat in optimal water conditions as shown by the size of the boxes in Fig. 7A. The leaves length of ryegrass and tall fescue was significantly longer at elevated [CO_2_]. Even if this study focused on the leaf growth response, our results also showed a strong effect of [CO_2_] on tillering. We calculated a tiller index (tillers (excluding the main stem) DM over the DM of the main stem), which increased at elevated [CO_2_] and mainly explained the increase in leaf area for both perennial grasses. Other studies such as Brinkhoff *et al*. (2018), Nicolas *et al*. (1984), Sionit *et al*. (1981) also reported that [CO_2_] increased tillering of ryegrass and wheat. As [CO_2_] had no significant impact on the rate of leaf emergence in all species (HI, Fig. 3B), it can be assumed that the differences in tillering between [CO_2_] = 200 and 800 ppm were not related to the number of lateral apices, which is co-ordinated with leaf initiation (Davies, 1974), but rather to the further differentiation of axillary apices in new tillers. In our experiment, it seems that elevated [CO_2_] increased the fraction of apices which ultimately develop into a visible tiller. In their study on herbaceous species, Ackerly *et al*. (1992) did not found any effect of CO_2_ on rates of leaf initiation and explained tillering with effects on the overall rate of development. At elevated CO_2_, in ryegrass, the lowest WSC accumulation while the highest tillering, suggest a greater WSC use for tillers production than the two other species. Baca Cabrera *et al*. (2020) did not observe an effect of [CO_2_] on WSC concentration in ryegrass and concluded that WSC was not limiting ryegrass developpement, but effect on tillering was not presented. The reduction of tillering at low [CO_2_] can be due to a lack of water and/or sugars for plant growth. Davies (1965) reported that site filling varies under environmental conditions and decreases with low carbohydrate availability. Our results showed a large difference of tillering and WSC content between 200 and 800 ppm in watered plants of wheat and tall fescue (Fig. 7A), which suggest a limitation by WSC.

CO_2_ effect on the relative variation of the morphogenesis variables was stronger during drought for all three species (Fig. 7B), especially for wheat which showed similar size boxes as ryegrass. Ryegrass presented the same type of response as in optimal conditions, with a very high tiller index linked to WSC use. Tiller index in ryegrass was stimulated by CO_2_ in much higher proportions than tall fescue and wheat. CO_2_ strongly enhanced the Specific Leaf Area (SLA) and leaf area of wheat plants (Fig. 7B), suggesting that CO_2_ mainly affected the width, thickness or composition rather than the length of the wheat leaves during drought. CO_2_ enhanced the leaf length and plant biomass of tall fescue to a greater extent than tillering during drought. The large increase in WSC content not invested into growth, probably explained the weaker relative response to CO_2_ of tall fescue compared to the one of the other species (Fig. 7B).

In addition to the analysis on shoot growth, we studied the development of underground parts and data showed that CO_2_ stimulated both roots and above-ground biomass (Fig. 7A). The extent of roots has implications for the efficiency of water and mineral extraction from soil to sustain shoot growth (Pritchard *et al*., 1999). CO_2_ did not affect the allocation of resources in response to drought (non-significant p-value, Supplementary Data Fig. S6).

## 5. CONCLUSION

In conclusion, our work enabled us to quantify the interaction of [CO_2_] and water availability on LER of individual plants for a long period during which several leaf series have been developed. This study revealed that [CO_2_] markedly mitigated and delayed the impacts of water deficit on LER of wheat, tall fescue and ryegrass. The positive effect of CO_2_ on LER was mainly due to the reduction of transpiration induced by stomatal closure. Interestingly, the reduction of water loss by plants did not affect water and osmotic potential response to [CO_2_] during nighttime, despite WSC accumulation. On the one hand, elevated [CO_2_] allowed preservation of water and prevention of water deficit damage in the early stages. On the other hand, leaf growth enhancement induced by elevated [CO_2_] resulted in similar (tall fescue and wheat) or higher (ryegrass) water consumption at elevated CO_2_. Leaf area increase was mainly due to the effect of CO_2_ on tillering.

The plant response to CO_2_ x drought interaction showed a large variability between species. Ryegrass was characterized by a rapid development rate and presented the lowest gs_w_ and instantaneous transpiration rate at the end of the experiment, as well as the lowest WSC accumulation in both sheaths and blades, while tillering was particularly high at [CO_2_] = 800 ppm. The results suggest that both WSC and water were invested into the growth of tillers of ryegrass. Tall fescue produced longest leaves, higher plant biomass and WSC at elevated [CO_2_] under drought, but showed a lower Haun Index and therefore a lower tillering activity than ryegrass. We observed a higher SLA, leaf area and WSC accumulation with CO_2_ in wheat that could be due to variety selection to foster grains rather than vegetative production. While perennial species responded in greater proportions to CO_2_ under optimal irrigation, CO_2_ stimulated wheat growth as much as ryegrass during drought whereas tall fescue was less responsive due to lower tillering. Building on the present interspecific variability among Poaceae, studies of the LER response to [CO_2_] x drought interaction should be expanded to multiple cultivars per species to further investigate the intraspecific variability of those results.

## SUPPLEMENTARY DATA

The following supplementary data are available at EEB online.

**Fig. S1.**
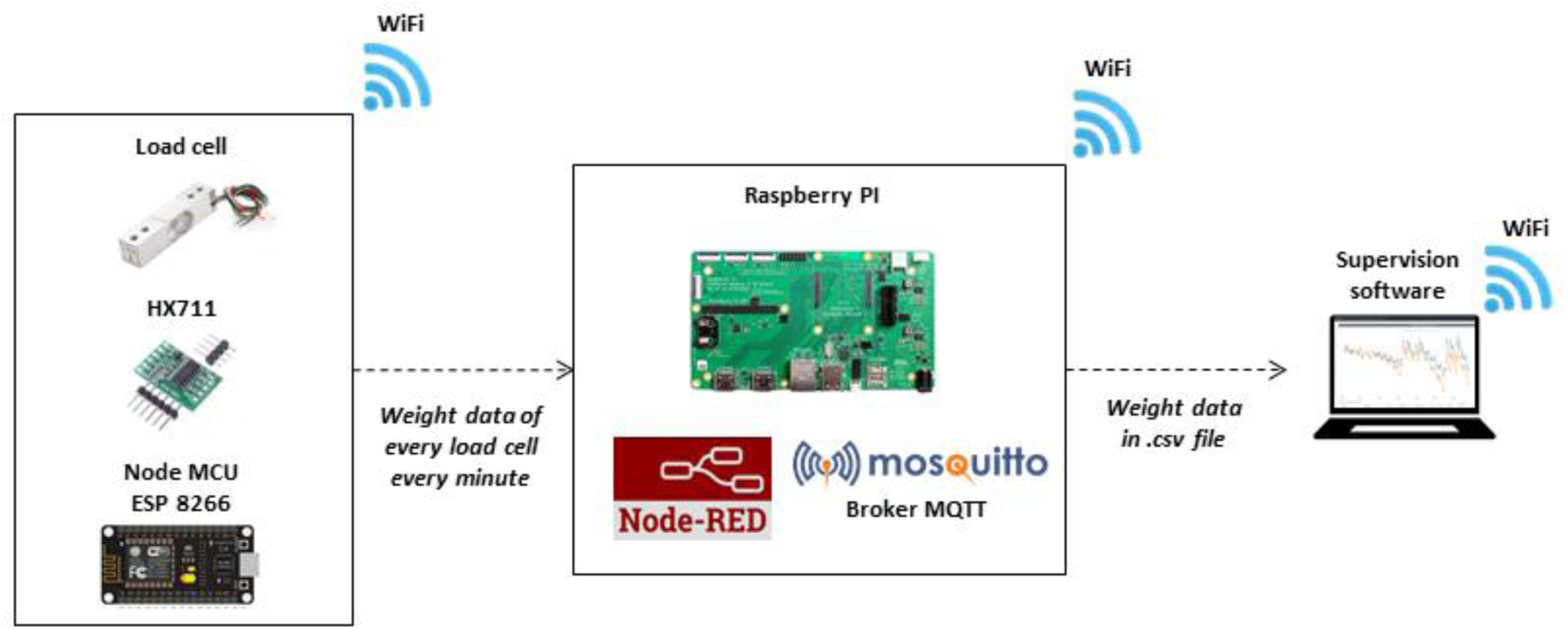
Explanatory diagram of the connection between load cells and the supervision software. *Each load cell (2 kg capacity) and its amplifier (HX711) was linked to a Node MCU card (ESP 8266) and connected to a WiFi network. The software Node-RED and the Broker MQTT Mosquitto were installed on a Raspberry PI connected to the same WiFi network than the load cells. Node-Red was programmed to request to data of each load cell every minute. Analog signal from load cell was transferred to the amplifier which translated it into a numerical signal for the Node MCU. Signal was transferred to Broker MQTT and then to Node-Red to process data and saved it in .csv file on the Raspberry PI memory card. The supervision software gathered the data of the .csv file and empty it. Node-Red was also programmed for plant water supply. Node MCU transferred the irrigation data via the Broker MQTT and the Node-Red to the supervision program in the same way that for weighting data*. *To overcome the vibration disturbances related to the functioning of the growth chambers, the load cells were fixed on an 8 mm thick aluminium plate. An insulator was placed on the top of each load cell to limit temperature variations*.

**Fig. S2.**
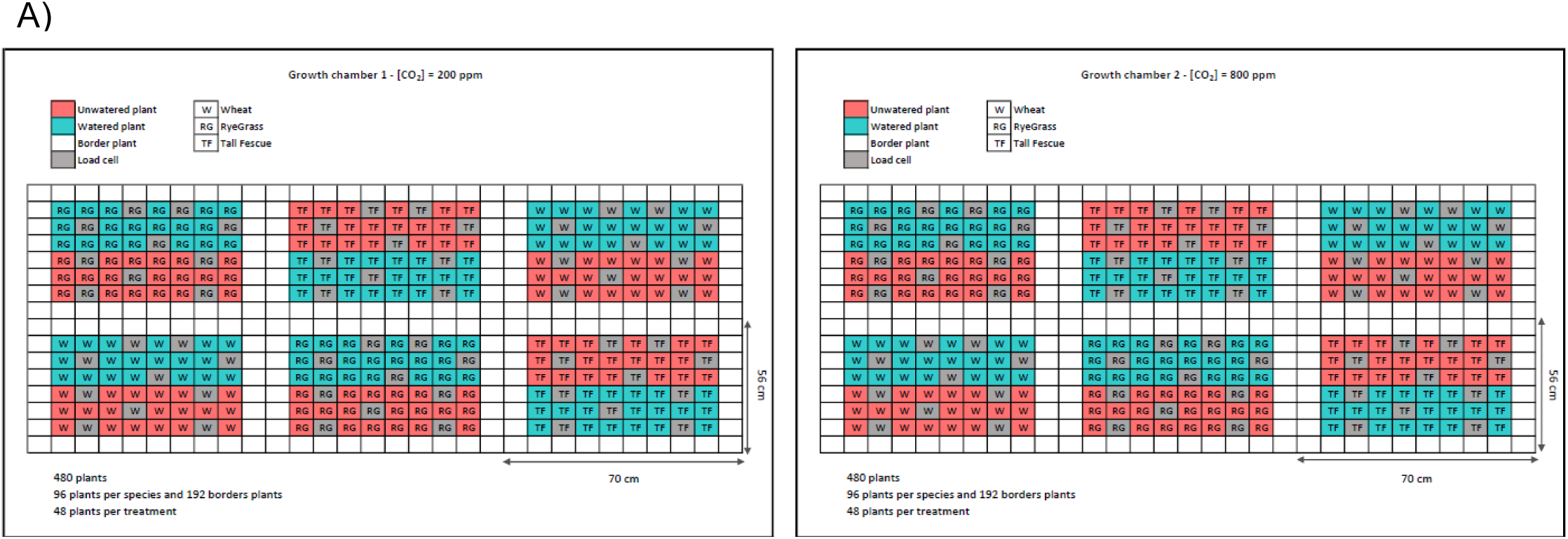
Experimental design of the growth chambers. *The growth chambers were set at [CO_2_] = 200 ppm (left side) and at [CO_2_] = 800 ppm (right side). Plants under watered treatment are represented in blue colour and unwatered treatment in red. Border plants are coloured in white and load cells in grey. The experiment was conducted on Wheat (Triticum aestivum L.; W*)*, RyeGrass (Lolium perenne L.; RG*) *by and Tall Fescue (Festuca arundinacea; TF*)*. Six blocks of plants (two per treatment) were disposed randomly in each chamber to limit the effect of spatial heterogeneity*.

**Table S1.**
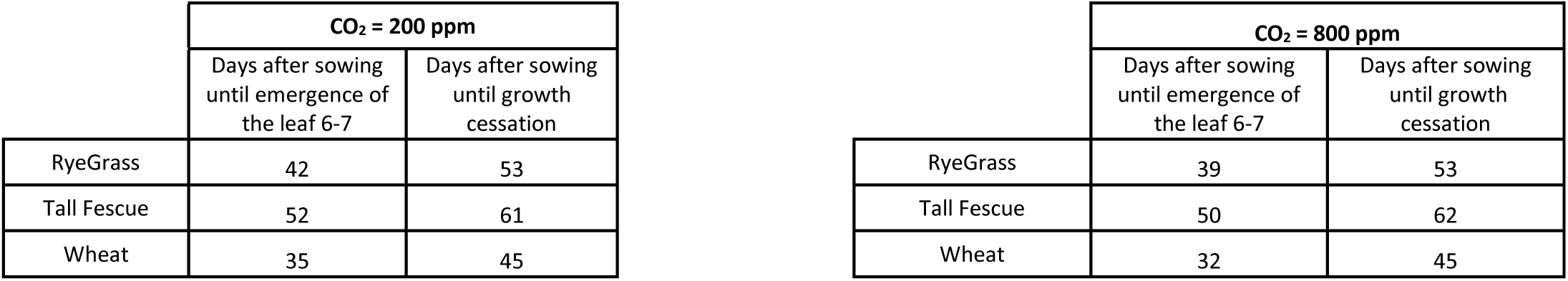
Timeline experiment for each treatment from sowing to emergence of leaf 6-7 (drought onset) and at to growth cessation for unwatered plants.

**Fig. S3.**
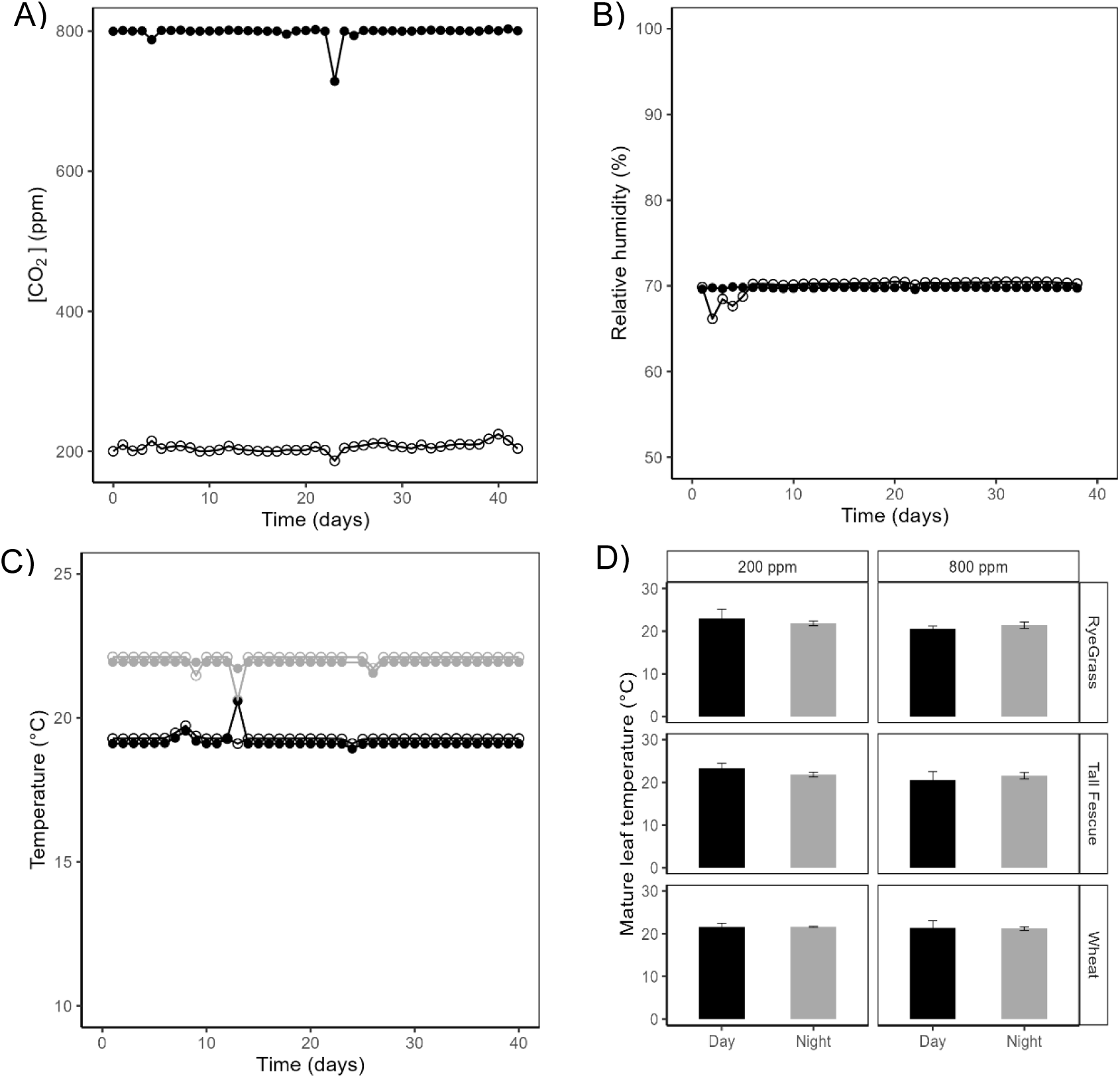
Daily mean of CO_2_ concentration (ppm) (A), relative humidity (%) (B), air temperature (°C) (C), and mature leaf temperature (°C) (D) in the two growth chambers. *Data are shown for plants grown at 200 (open circles) or 800 ppm [CO_2_] (closed circles) during daytime (black) or nighttime (grey) conditions*. *PAR was measured at the vegetation height in each chamber by scanning the entire surface in 15 points, showing constant radiation between CO_2_ treatments (*∼*442.6 ± 11.2 µmol PAR m^-2^ s^-1^ at 200 ppm and* ∼*414.7 ± 29.5 µmol PAR m^-2^ s^-1^ at 800 ppm)*.

**Fig. S4.**
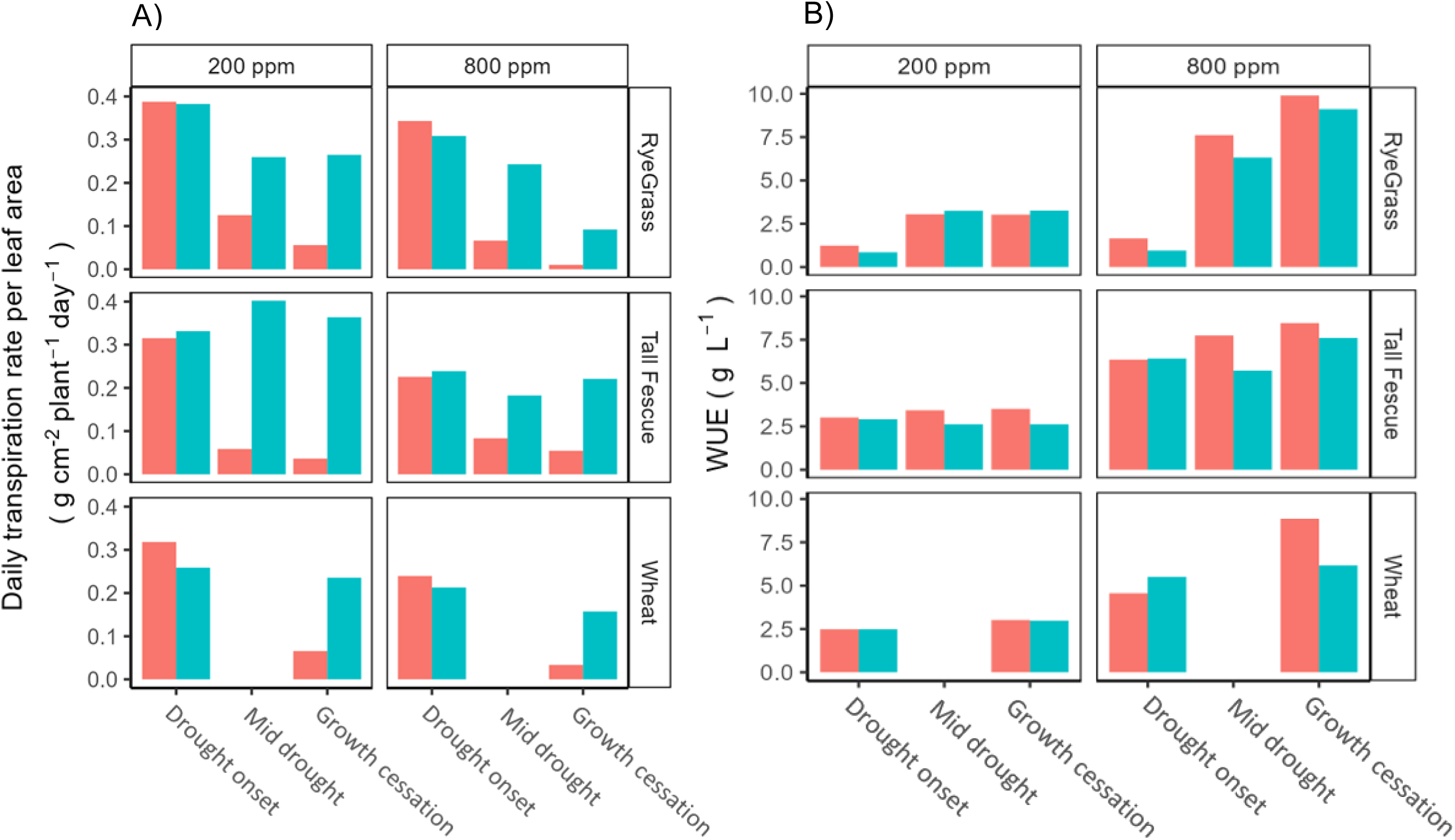
Transpiration rate per unit of leaf area (g H_2_O cm^-2^ plant^-1^) (A) and water-use efficiency (WUE, g L^-1^) (A) for *Lolium perenne L.* (upper panels), *Festuca arundinacea* (mid-panels) and *Triticum aestivum L.* (bottom panels) at three drought steps: drought onset, mid-drought and growth cessation. *Results are shown for plants grown at 200 (left side panels) or 800 ppm [CO_2_] (right side panels) in watered (blue) or unwatered (red) conditions. Data were averaged on five plants per treatment. WUE was calculated as the ratio between plant dry matter (g plant^-1^) and transpiration (L plant^-1^)*.

**Fig. S5.**
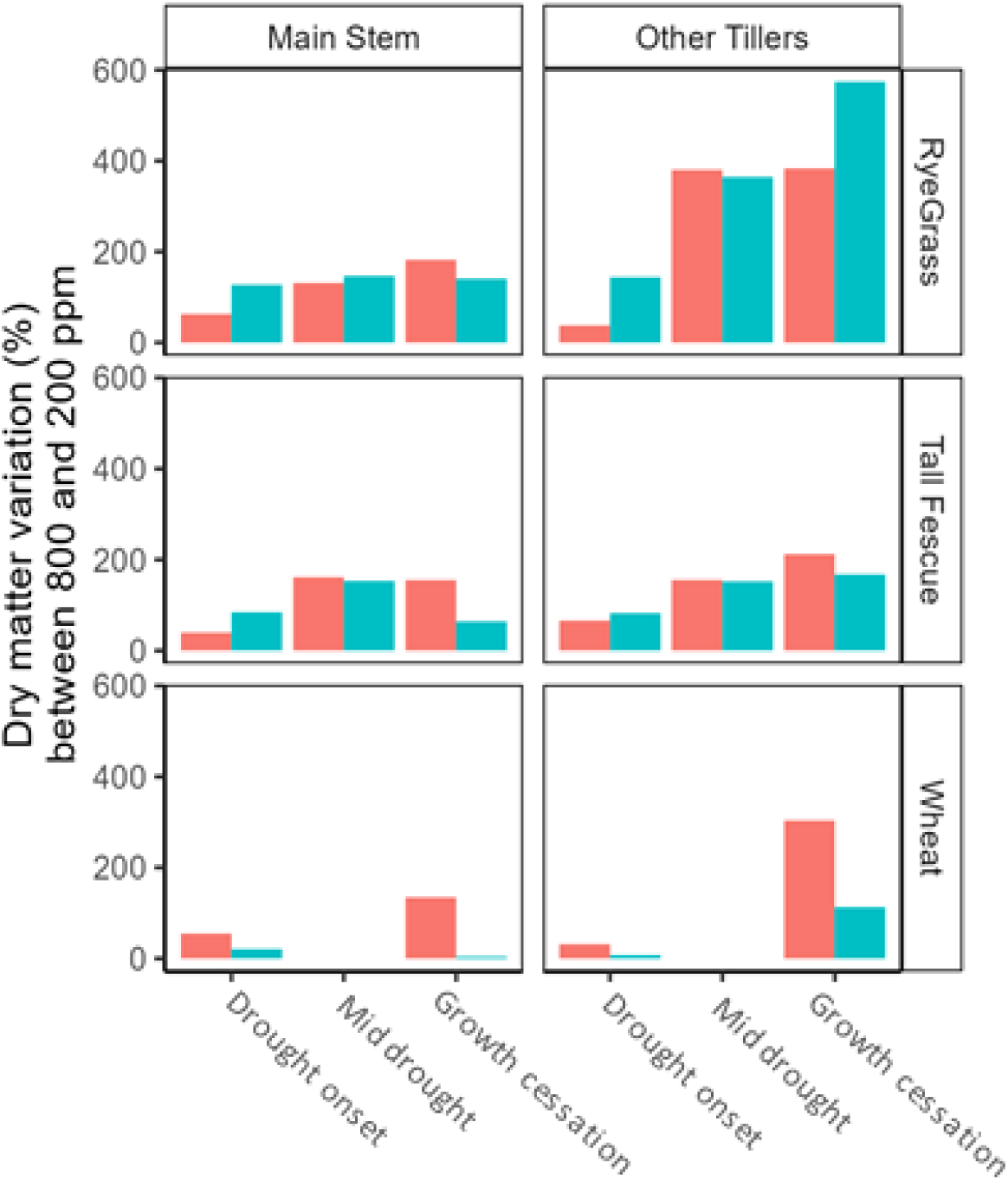
Dry matter variation (%) of main stem (left panels) and other tillers (right panels) between plants grown at 200 or 800 ppm of CO_2_ for *Lolium perenne L.* (upper panels), *Festuca arundinacea* (mid-panels) and *Triticum aestivum L.* (bottom panels) at three drought step: drought onset, mid-drought and growth cessation. *Results are shown for plants grown at 200 (open circles) or 800 ppm [CO_2_] (closed circles) in watered (blue) or unwatered (red) conditions. Data is averaged on five plants per treatment*.

**Fig. S6.**
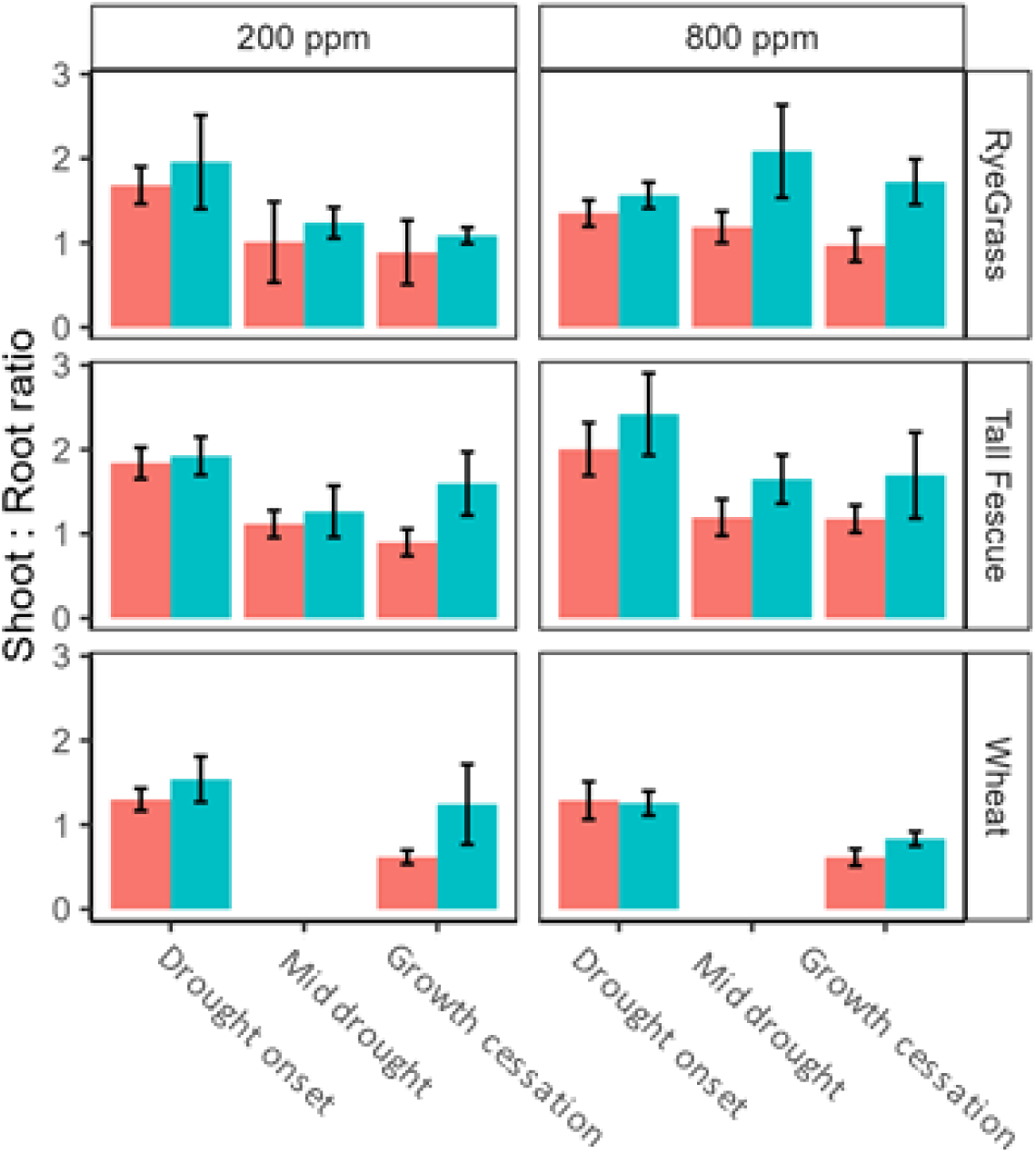
Shoot: Root ratio for *Lolium perenne L.* (upper panels), *Festuca arundinacea* (mid-panels) and *Triticum aestivum L.* (bottom panels) at three drought step: drought onset, mid-drought and growth cessation. *Results are shown for plants grown at 200 (left side panels) or 800 ppm [CO_2_] (right side panels) in watered (blue) or unwatered (red) conditions. Data is averaged on five plants per treatment. Error bars represent the mean ± standard deviation*.

**Table S2.**
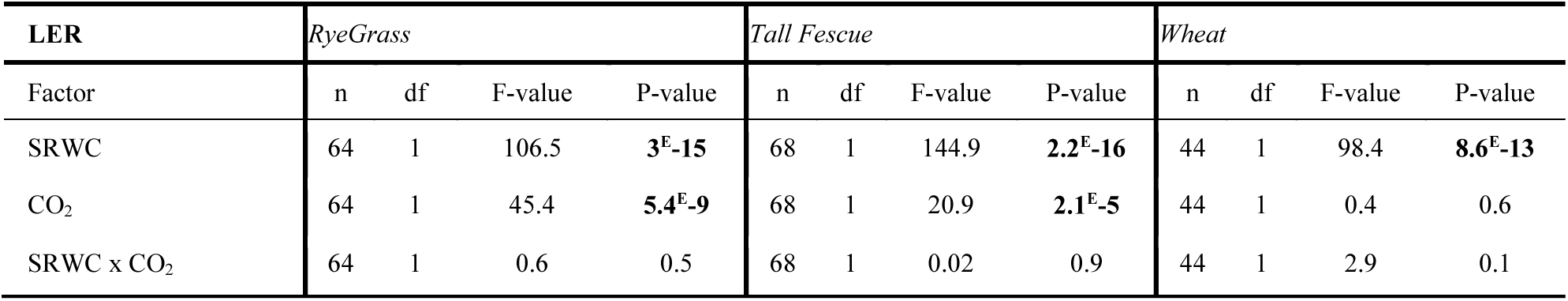
Results of a linear mixed model, testing the response of daily mean leaf elongation rate to soil relative water content (SRWC, %), atmospheric CO_2_ concentration ([CO_2_]) (800 μmol mol^−1^ vs 200 μmol mol^−1^), and their interaction for *Lolium perenne L*., *Festuca arundinacea* and *Triticum aestivum L*.

**Table S3.**
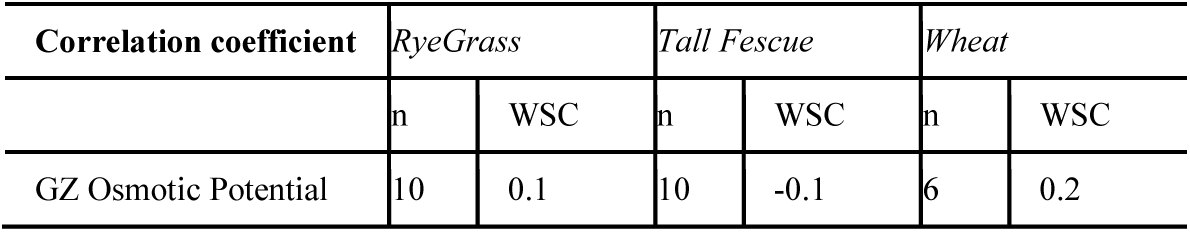
Correlation coefficients of the relation between the growing zone osmotic potential (π _GZ_, Mpa) and water-soluble carbohydrates (WSC, % dry matter) or *Lolium perenne L*., *Festuca arundinacea* and *Triticum aestivum L*.

| Correlation coefficient | *RyeGrass* |  | *Tall Fescue* |  | *Wheat* |  |
| --- | --- | --- | --- | --- | --- | --- |
|  | n | WSC | n | WSC | n | WSC |
| GZ Osmotic Potential | 10 | 0.1 | 10 | -0.1 | 6 | 0.2 |

**Table S4.**
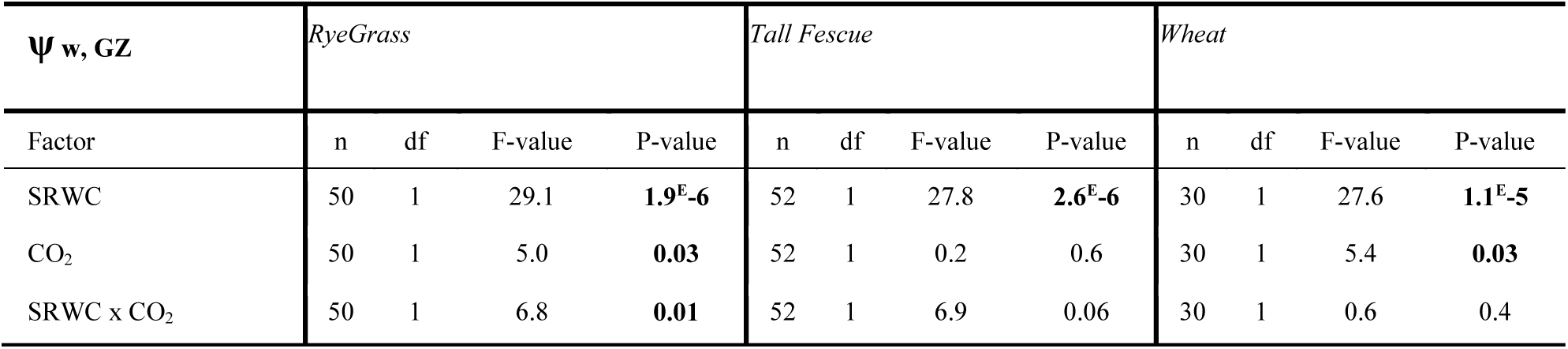
Results of a linear mixed model, testing the response of GZ water potential (ψ_GZ_, Mpa) to soil relative water content (SRWC, %), atmospheric CO_2_ concentration ([CO_2_]) (800 μmol mol^−1^ vs 200 μmol mol^−1^), and their interaction for *Lolium perenne L*., *Festuca arundinacea* and *Triticum aestivum L*.

**Fig. S7.**
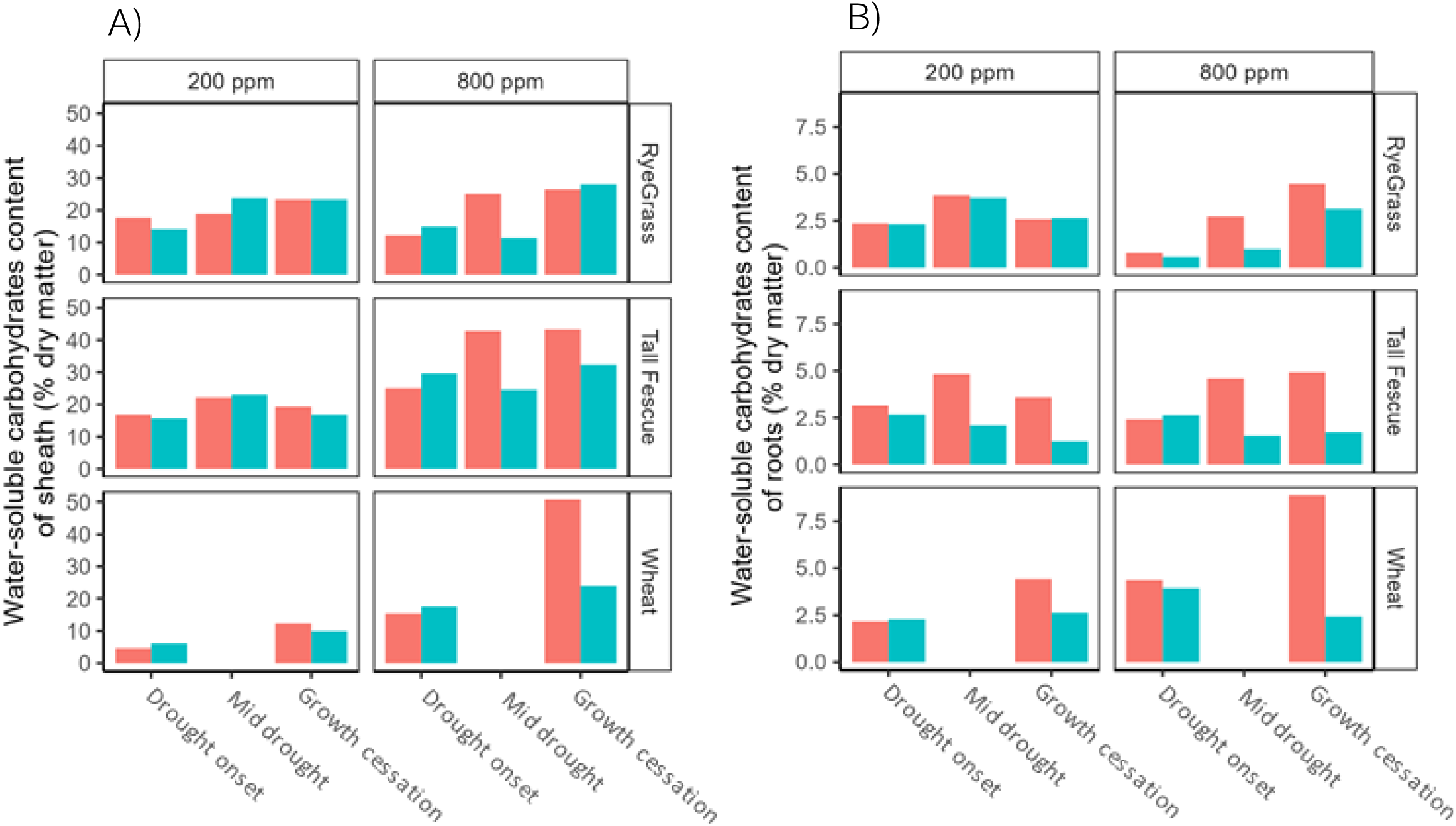
Water-soluble carbohydrate content (% dry matter plant) of sheath (A) and roots (B) for *Lolium perenne L.* (upper panels), *Festuca arundinacea* (mid-panels) and *Triticum aestivum L.* (bottom panels) at three drought stages: drought onset, mid-drought and growth cessation. *Results are shown for plants grown at 200 (left side panel) or 800 ppm [CO_2_] (right side panel) in watered (blue) or unwatered (red) conditions. Data is averaged on five plants per treatment*.

**Table S5.**
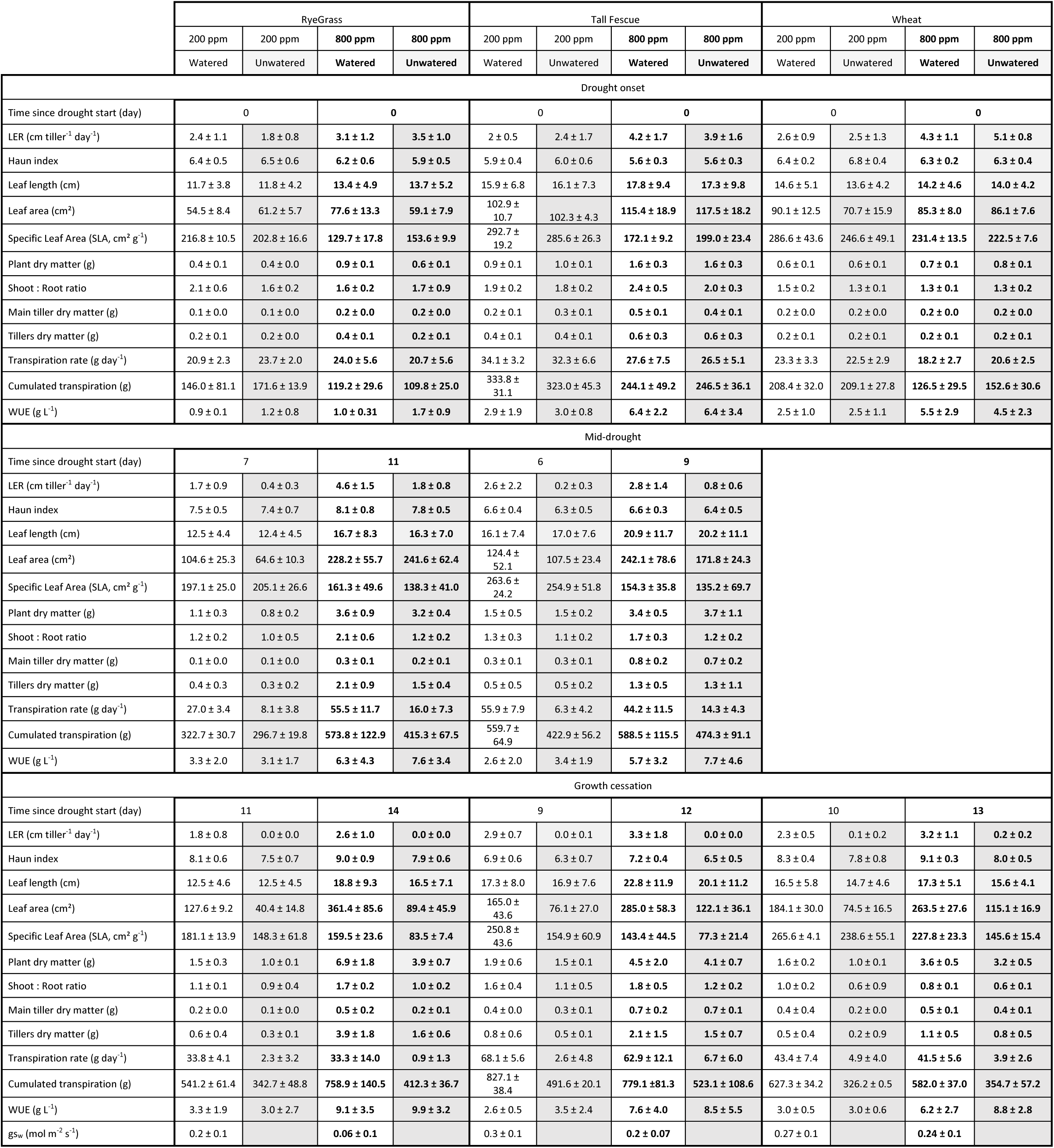
Mean ± standard deviation data of 10 plants per treatment of Leaf Elongation Rate (LER) (cm tiller^-1^ day^-1^), leaf length (cm), Haun Index, plant dry matter (g), Shoot: Root ratio, main stem dry matter (g), tiller dry matter (g), leaf area (cm²), Specific Leaf Area (SLA) (cm² g^-1^), transpiration rate (g day^-1^),cumulated transpiration (g), and WUE (g L^-1^) for *Lolium perenne L.*, *Festuca arundinacea* and *Triticum aestivum L.,* at three drought step: drought onset, mid-drought, growth cessation.

**Table S6.**
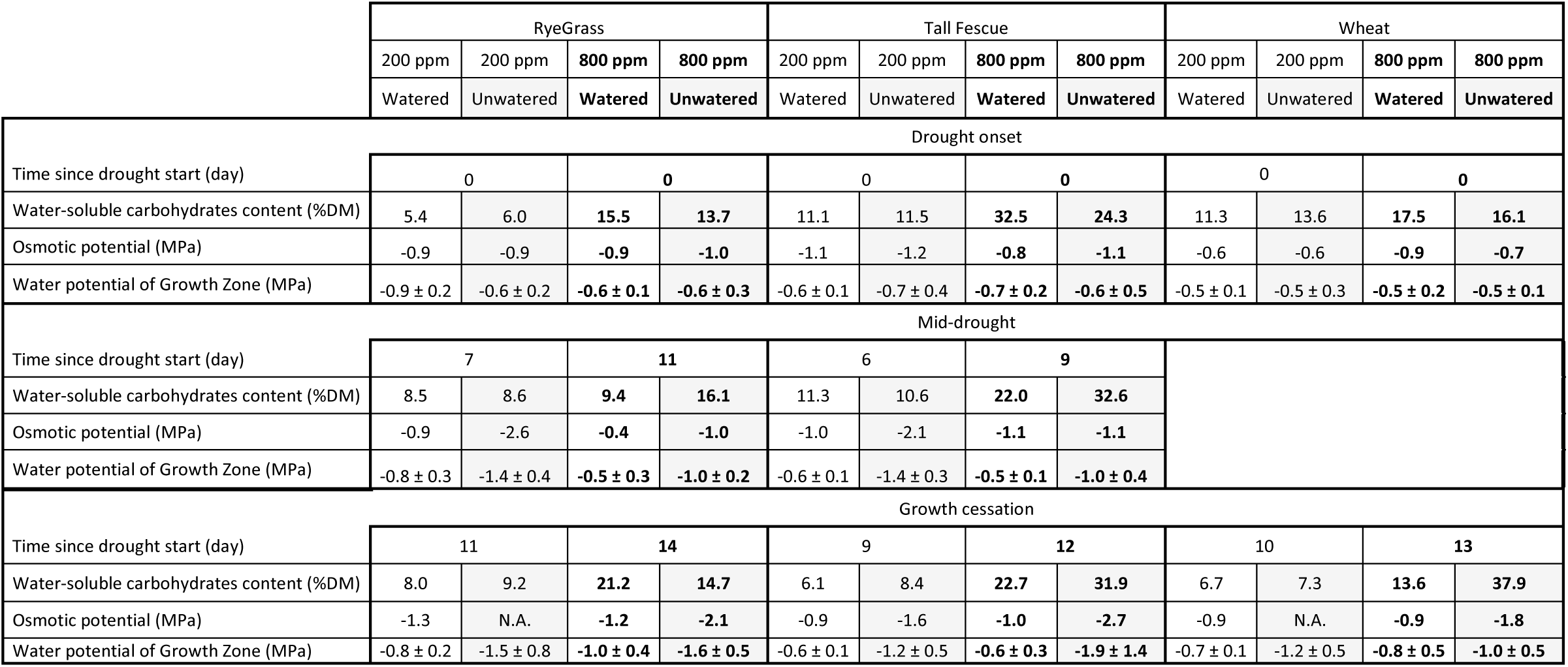
Mean ± standard deviation data of 10 plants per treatment of water-soluble carbohydrates (% dry matter), osmotic potential (MPa), turgor potential (MPa), water potential of mature leaves and growing zone (MPa), potential gradient of mature leaf and growing zone (MPa), for *Lolium perenne L.*, *Festuca arundinacea* and *Triticum aestivum L.,* at three drought step: drought onset, mid-drought, growth cessation.

## ACKNOWLEDGEMENTS

The authors thank Franck Gelin, Pascal Vernoux, Annie Eprinchard, Marianne van Peteghem and Nathan Merle for their technical assistance.

## AUTHOR CONTRIBUTIONS

The experiment was designed by RB. The growth chambers with devices to control the CO_2_ and water treatments, as well as the load cell system below the plants, were installed by CP and ER. The acquisition, analysis, and interpretation of the data was performed by VA. Gas exchange was measured by EF. VA, RB and JLD contributed to the review of the literature, conceptualisation of thinking, and writing.

## CONFLICT OF INTERESTS

No conflict of interest declared.

## FUNDING

This work was supported by a convention between the Région Nouvelle Aquitaine and AgroEcoSystèmes Department of Institut national de recherche pour l’agriculture, l’alimentation et l’environnement (INRAe) under V.A.’s PhD (Convention n° AAPR2021A-2020-11767810).

## DATA AVAILABILITY

Numerical data used in the figures are available in Tables S1 and S2. Raw data are available upon request from the corresponding authors.

## Notes

### Competing Interest Statement

The authors have declared no competing interest.

