## Supplementary Data for "Effects of atmospheric CO_2_ concentration on transpiration and leaf elongation responses to drought in wheat, perennial ryegrass and tall fescue"

**Victoria Acker[1,*], Jean-Louis Durand[1], Cédric Perrot[1], Eric Roy[1], Elzbieta Frak[1] and Romain Barillot[1,*]**
[1]INRAE, UR P3F, F-86600 Lusignan, France

### SUPPLEMENTARY DATA

The following supplementary data are available online.

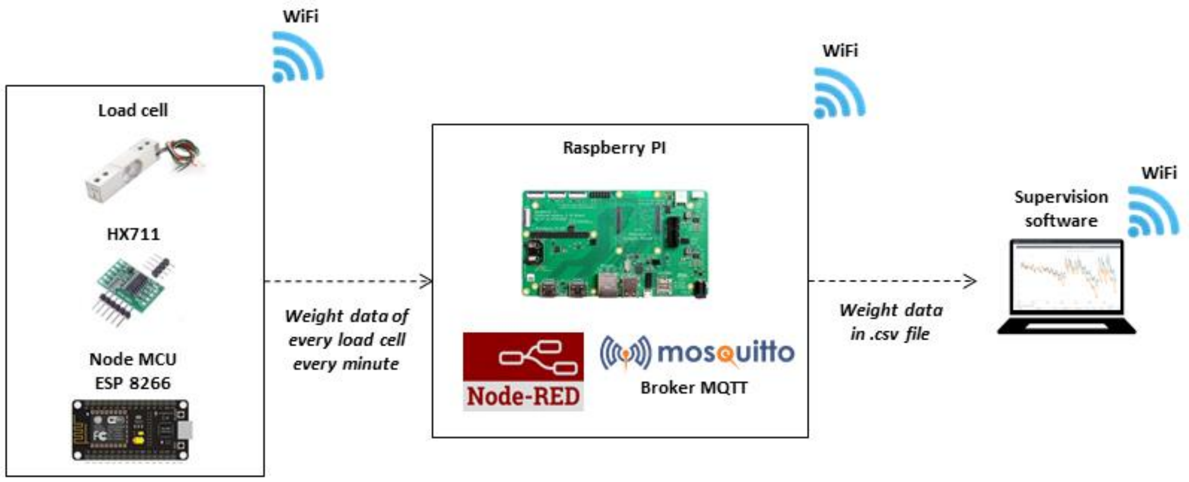


**Fig. S1. Explanatory diagram of the connection between load cells and the supervision software.**

*Each load cell (2 kg capacity) and its amplifier (HX711) was linked to a Node MCU card (ESP 8266) and connected to a WiFi network. The software Node-RED and the Broker MQTT Mosquitto were installed on a Raspberry PI connected to the same WiFi network than the load cells. Node-Red was programmed to request to data of each load cell every minute. Analog signal from load cell was transferred to the amplifier which translated it into a numerical signal for the Node MCU. Signal was transferred to Broker MQTT and then to Node-Red to process data and saved it in .csv file on the Raspberry PI memory card. The supervision software gathered the data of the .csv file and empty it. Node-Red was also programmed for plant water supply. Node MCU transferred the irrigation data via the Broker MQTT and the Node-Red to the supervision program in the same way that for weighting data.*
*To overcome the vibration disturbances related to the functioning of the growth chambers, the load cells were fixed on an 8 mm thick aluminium plate. An insulator was placed on the top of each load cell to limit temperature variations.*

A)

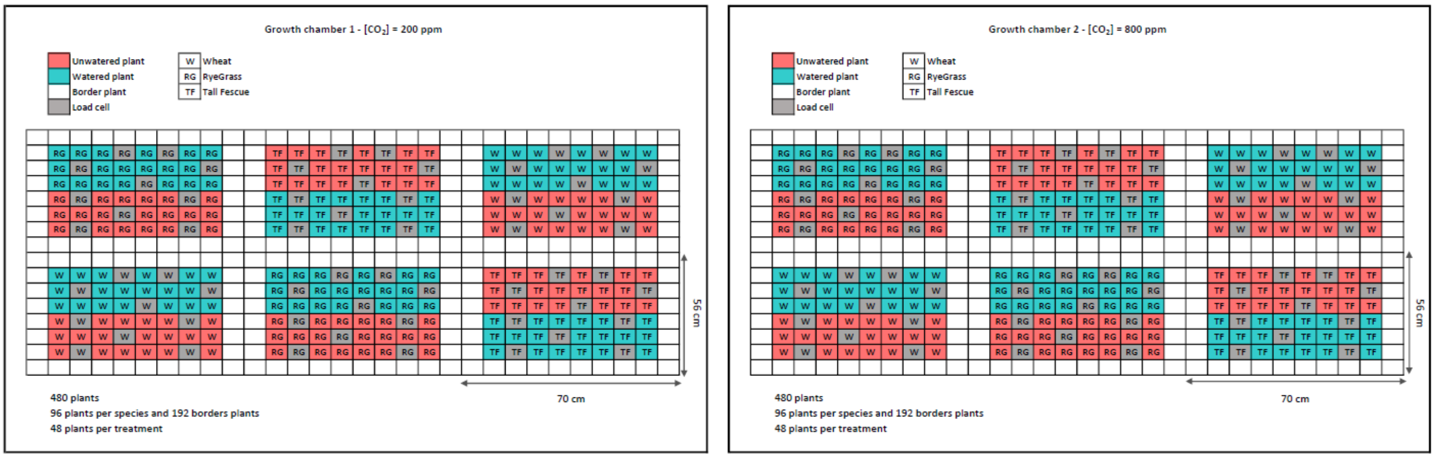


**Fig. S2. Experimental design of the growth chambers.**

*The growth chambers were set at [$CO_2$] = 200 ppm (left side) and at [$CO_2$] = 800 ppm (right side). Plants under watered treatment are represented in blue colour and unwatered treatment in red. Border plants are coloured in white and load cells in grey. The experiment was conducted on Wheat (Triticum aestivum L. ; W), RyeGrass (Lolium perenne L. ; RG) by and Tall Fescue (Festuca arundinacea ; TF). Six blocks of plants (two per treatment) were disposed randomly in each chamber to limit the effect of spatial heterogeneity.*

**Table S1. Timeline experiment for each treatment from sowing to emergence of leaf 6-7 (drought onset) and at to growth cessation for unwatered plants.**

| | $CO_2$ = 200 ppm | |
|---|---|---|
| | Days after sowing until emergence of the leaf 6-7 | Days after sowing until growth cessation |
| RyeGrass | 42 | 53 |
| Tall Fescue | 52 | 61 |
| Wheat | 35 | 45 |

| | $CO_2$ = 800 ppm | |
|---|---|---|
| | Days after sowing until emergence of the leaf 6-7 | Days after sowing until growth cessation |
| RyeGrass | 39 | 53 |
| Tall Fescue | 50 | 62 |
| Wheat | 32 | 45 |

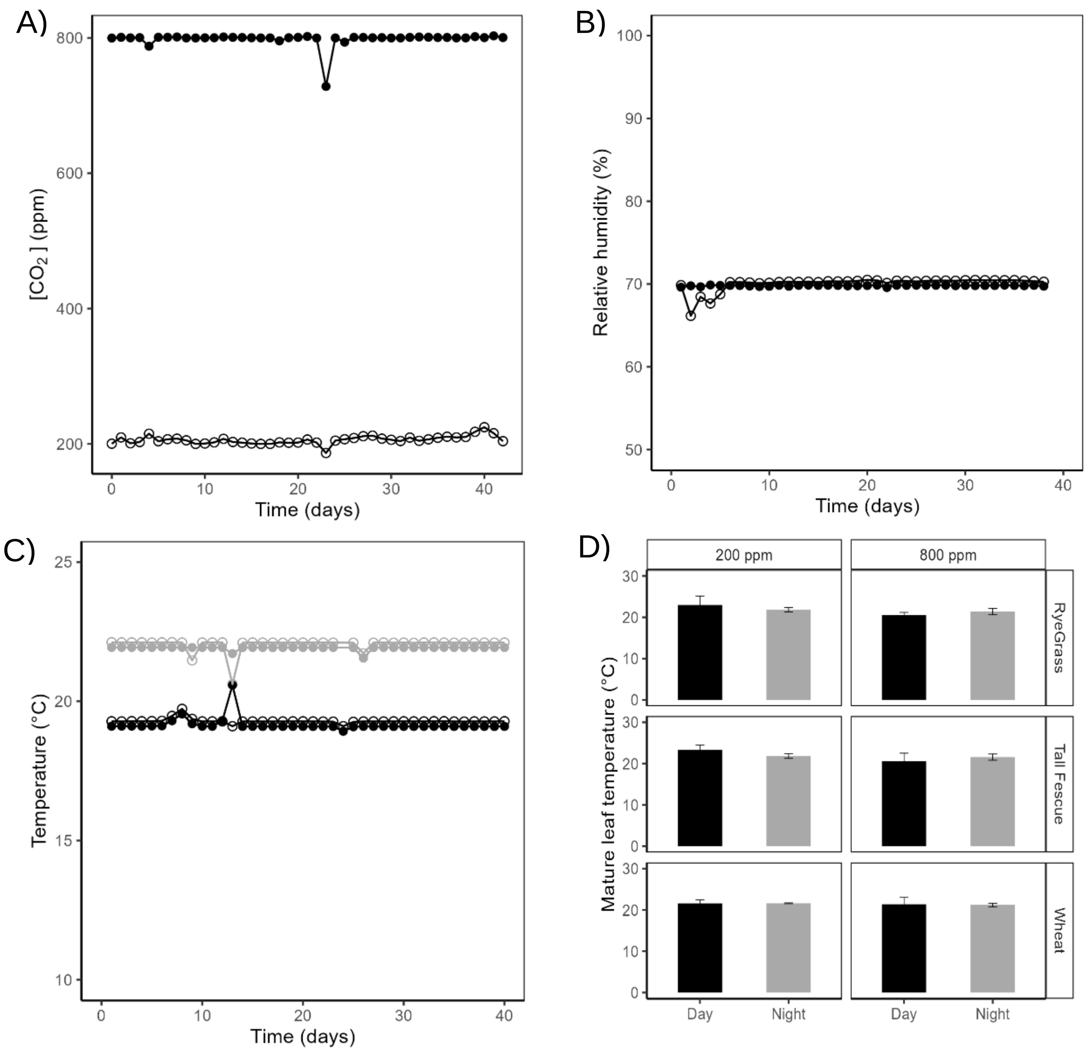


**Fig. S3. Daily mean of $CO_2$ concentration (ppm) (A), relative humidity (%) (B), air temperature (°C) (C), and mature leaf temperature (°C) (D) in the two growth chambers.**

*Data are shown for plants grown at 200 (open circles) or 800 ppm [$CO_2$] (closed circles) during daytime (black) or nighttime (grey) conditions.*

*PAR was measured at the vegetation height in each chamber by scanning the entire surface in 15 points, showing constant radiation between $CO_2$ treatments (~442.6 ± 11.2 µmol PAR $m^{-2}$ $s^{-1}$ at 200 ppm and ~414.7 ± 29.5 µmol PAR $m^{-2}$ $s^{-1}$ at 800 ppm).*

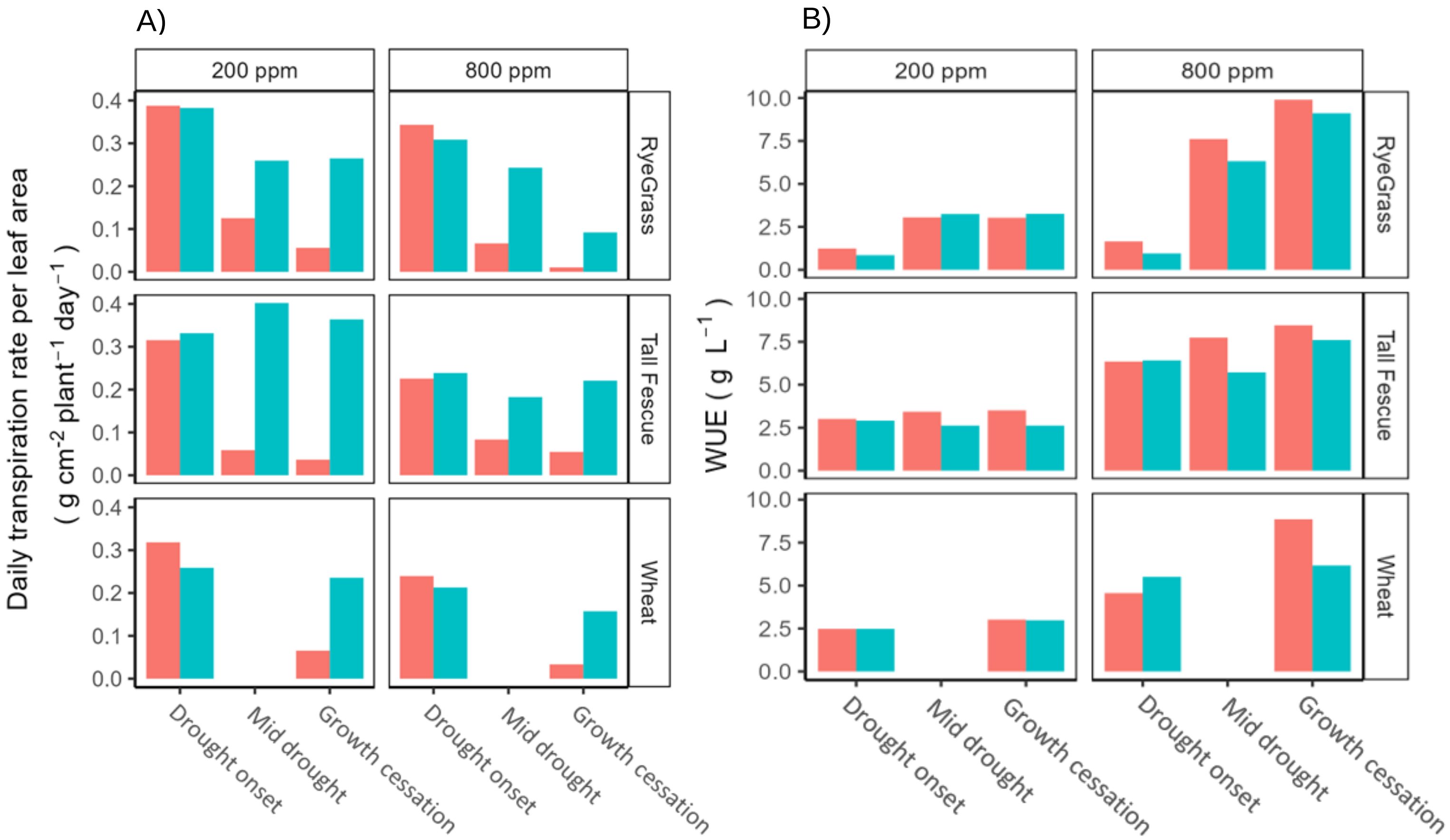


**Fig. S4. Transpiration rate per unit of leaf area (g $H_2O$ $cm^{-2}$ $plant^{-1}$) (A) and water-use efficiency (WUE, g $L^{-1}$) (A) for *Lolium perenne L.* (upper panels), *Festuca arundinacea* (mid-panels) and *Triticum aestivum L.* (bottom panels) at three drought steps: drought onset, mid-drought and growth cessation.**

*Results are shown for plants grown at 200 (left side panels) or 800 ppm [$CO_2$] (right side panels) in watered (blue) or unwatered (red) conditions. Data were averaged on five plants per treatment. WUE was calculated as the ratio between plant dry matter (g $plant^{-1}$) and transpiration (L $plant^{-1}$).*

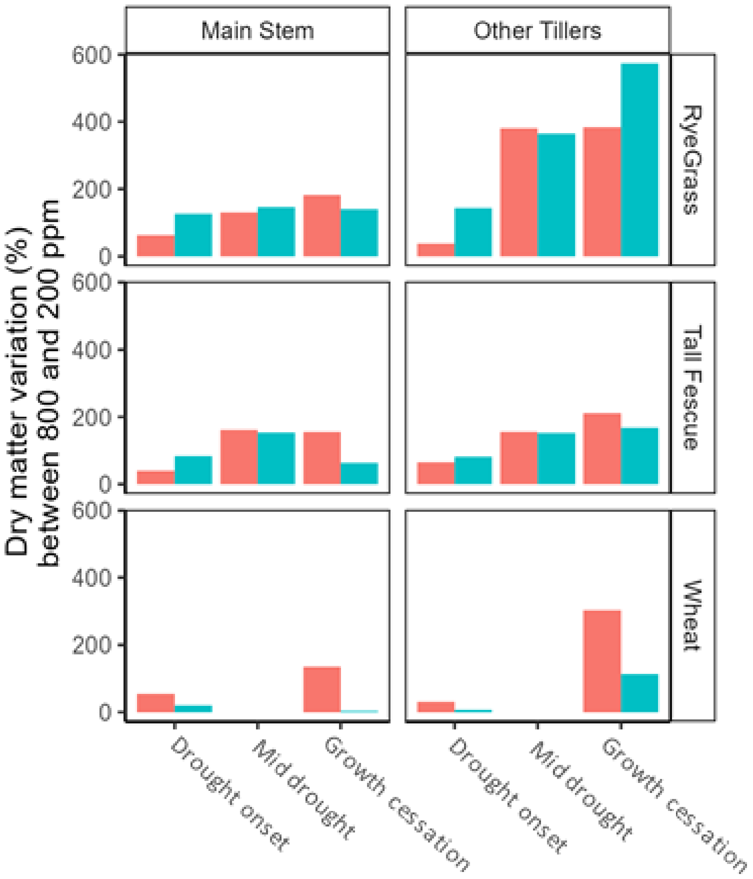


**Fig. S5. Dry matter variation (%) of main stem (left panels) and other tillers (right panels) between plants grown at 200 or 800 ppm of $CO_2$ for *Lolium perenne L.* (upper panels), *Festuca arundinacea* (mid-panels) and *Triticum aestivum L.* (bottom panels) at three drought step: drought onset, mid-drought and growth cessation.**

*Results are shown for plants grown at 200 (open circles) or 800 ppm [$CO_2$] (closed circles) in watered (blue) or unwatered (red) conditions. Data is averaged on five plants per treatment.*

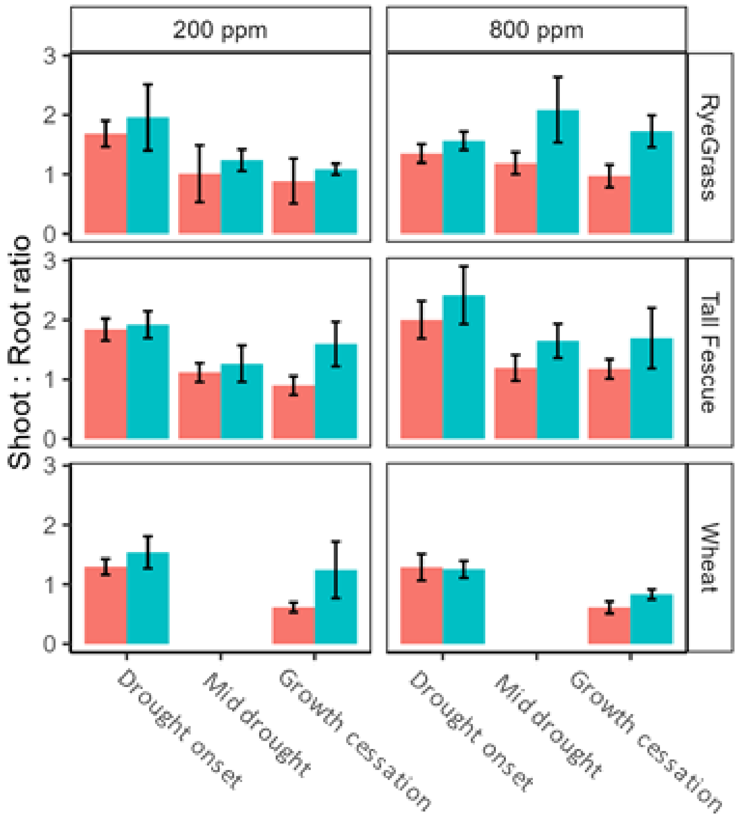


**Fig. S6. Shoot : Root ratio for *Lolium perenne L.* (upper panels), *Festuca arundinacea* (mid-panels) and *Triticum aestivum L.* (bottom panels) at three drought step: drought onset, mid-drought and growth cessation.**

*Results are shown for plants grown at 200 (left side panels) or 800 ppm [$CO_2$] (right side panels) in watered (blue) or unwatered (red) conditions. Data is averaged on five plants per treatment. Error bars represent the mean ± standard deviation.*

**Table S2. Results of a linear mixed model, testing the response of daily mean leaf elongation rate to soil relative water content (SRWC, %), atmospheric $CO_2$ concentration ($[CO_2]$) (800 µmol $mol^{-1}$ vs 200 µmol $mol^{-1}$), and their interaction for *Lolium perenne L.*, *Festuca arundinacea* and *Triticum aestivum L.***

| **LER** | *RyeGrass* | | | | *Tall Fescue* | | | | *Wheat* | | | |
|---|---|---|---|---|---|---|---|---|---|---|---|---|
| Factor | n | df | F-value | P-value | n | df | F-value | P-value | n | df | F-value | P-value |
| SRWC | 64 | 1 | 106.5 | **3E-15** | 68 | 1 | 144.9 | **2.2E-16** | 44 | 1 | 98.4 | **8.6E-13** |
| $CO_2$ | 64 | 1 | 45.4 | **5.4E-9** | 68 | 1 | 20.9 | **2.1E-5** | 44 | 1 | 0.4 | 0.6 |
| SRWC x $CO_2$ | 64 | 1 | 0.6 | 0.5 | 68 | 1 | 0.02 | 0.9 | 44 | 1 | 2.9 | 0.1 |

**Table S3. Correlation coefficients of the relation between the growing zone osmotic potential ($\pi_{GZ}$, Mpa) and water-soluble carbohydrates (WSC, % dry matter) or *Lolium perenne L.*, *Festuca arundinacea* and *Triticum aestivum L.***

| **Correlation coefficient** | *RyeGrass* | | *Tall Fescue* | | *Wheat* | |
|---|---|---|---|---|---|---|
| | n | WSC | n | WSC | n | WSC |
| GZ Osmotic Potential | 10 | 0.1 | 10 | -0.1 | 6 | 0.2 |

**Table S4. Results of a linear mixed model, testing the response of GZ water potential ($\psi_{GZ}$, Mpa) to soil relative water content (SRWC, %), atmospheric $CO_2$ concentration ($[CO_2]$) (800 µmol mol$^{-1}$ vs 200 µmol mol$^{-1}$), and their interaction for *Lolium perenne L.*, *Festuca arundinacea* and *Triticum aestivum L.***

| **ψ w, GZ** | *RyeGrass* | | | | *Tall Fescue* | | | | *Wheat* | | | |
|---|---|---|---|---|---|---|---|---|---|---|---|---|
| Factor | n | df | F-value | P-value | n | df | F-value | P-value | n | df | F-value | P-value |
| SRWC | 50 | 1 | 29.1 | **1.9$^{E}$-6** | 52 | 1 | 27.8 | **2.6$^{E}$-6** | 30 | 1 | 27.6 | **1.1$^{E}$-5** |
| $CO_2$ | 50 | 1 | 5.0 | **0.03** | 52 | 1 | 0.2 | 0.6 | 30 | 1 | 5.4 | **0.03** |
| SRWC x $CO_2$ | 50 | 1 | 6.8 | **0.01** | 52 | 1 | 6.9 | 0.06 | 30 | 1 | 0.6 | 0.4 |

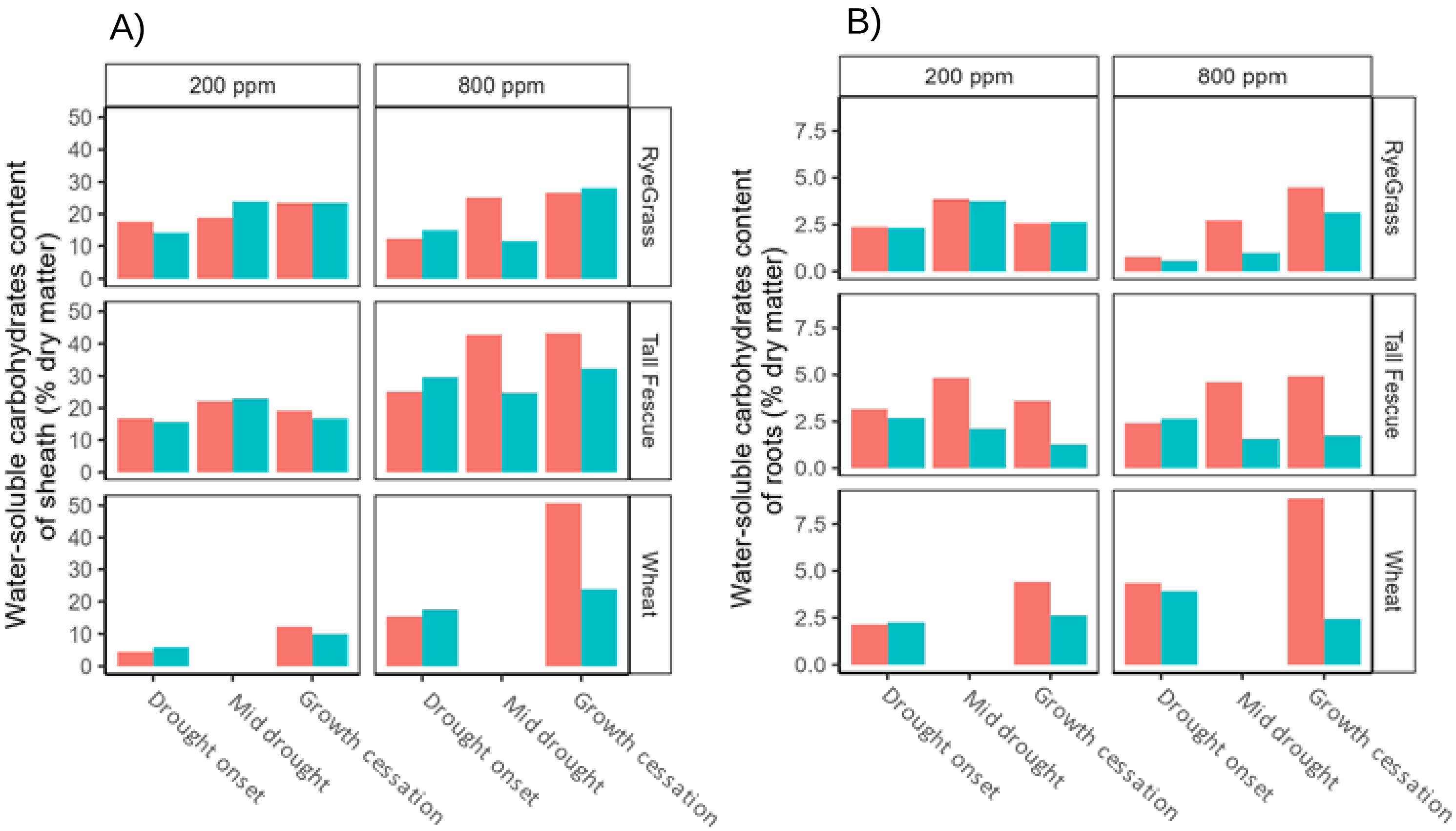


**Fig. S7. Water-soluble carbohydrate content (% dry matter plant) of sheath (A) and roots (B) for *Lolium perenne L.* (upper panels), *Festuca arundinacea* (mid-panels) and *Triticum aestivum L.* (bottom panels) at three drought stages: drought onset, mid-drought and growth cessation.**

*Results are shown for plants grown at 200 (left side panel) or 800 ppm [$CO_2$] (right side panel) in watered (blue) or unwatered (red) conditions. Data is averaged on five plants per treatment.*

**Table S5. Mean ± standard deviation data of 10 plants per treatment of Leaf Elongation Rate (LER) (cm tiller$^{-1}$ day$^{-1}$), leaf length (cm), Haun Index, plant dry matter (g), Shoot : Root ratio, main stem dry matter (g), tiller dry matter (g), leaf area (cm²), Specific Leaf Area (SLA) (cm² g$^{-1}$), transpiration rate (g day$^{-1}$),cumulated transpiration (g), and WUE (g L$^{-1}$) for *Lolium perenne L.*, *Festuca arundinacea* and *Triticum aestivum L.*, at three drought step: drought onset, mid-drought, growth cessation.**

| | RyeGrass | | | | Tall Fescue | | | | Wheat | | | |
|---|---|---|---|---|---|---|---|---|---|---|---|---|
| | 200 ppm | 200 ppm | **800 ppm** | **800 ppm** | 200 ppm | 200 ppm | **800 ppm** | **800 ppm** | 200 ppm | 200 ppm | **800 ppm** | **800 ppm** |
| | Watered | Unwatered | **Watered** | **Unwatered** | Watered | Unwatered | **Watered** | **Unwatered** | Watered | Unwatered | **Watered** | **Unwatered** |
| Drought onset | | | | | | | | | | | | |
| Time since drought start (day) | 0 | | **0** | | 0 | | **0** | | 0 | | **0** | |
| LER (cm tiller$^{-1}$ day$^{-1}$) | 2.4 ± 1.1 | 1.8 ± 0.8 | **3.1 ± 1.2** | **3.5 ± 1.0** | 2 ± 0.5 | 2.4 ± 1.7 | **4.2 ± 1.7** | **3.9 ± 1.6** | 2.6 ± 0.9 | 2.5 ± 1.3 | **4.3 ± 1.1** | **5.1 ± 0.8** |
| Haun index | 6.4 ± 0.5 | 6.5 ± 0.6 | **6.2 ± 0.6** | **5.9 ± 0.5** | 5.9 ± 0.4 | 6.0 ± 0.6 | **5.6 ± 0.3** | **5.6 ± 0.3** | 6.4 ± 0.2 | 6.8 ± 0.4 | **6.3 ± 0.2** | **6.3 ± 0.4** |
| Leaf length (cm) | 11.7 ± 3.8 | 11.8 ± 4.2 | **13.4 ± 4.9** | **13.7 ± 5.2** | 15.9 ± 6.8 | 16.1 ± 7.3 | **17.8 ± 9.4** | **17.3 ± 9.8** | 14.6 ± 5.1 | 13.6 ± 4.2 | **14.2 ± 4.6** | **14.0 ± 4.2** |
| Leaf area (cm²) | 54.5 ± 8.4 | 61.2 ± 5.7 | **77.6 ± 13.3** | **59.1 ± 7.9** | 102.9 ± 10.7 | 102.3 ± 4.3 | **115.4 ± 18.9** | **117.5 ± 18.2** | 90.1 ± 12.5 | 70.7 ± 15.9 | **85.3 ± 8.0** | **86.1 ± 7.6** |
| Specific Leaf Area (SLA, cm² g$^{-1}$) | 216.8 ± 10.5 | 202.8 ± 16.6 | **129.7 ± 17.8** | **153.6 ± 9.9** | 292.7 ± 19.2 | 285.6 ± 26.3 | **172.1 ± 9.2** | **199.0 ± 23.4** | 286.6 ± 43.6 | 246.6 ± 49.1 | **231.4 ± 13.5** | **222.5 ± 7.6** |
| Plant dry matter (g) | 0.4 ± 0.1 | 0.4 ± 0.0 | **0.9 ± 0.1** | **0.6 ± 0.1** | 0.9 ± 0.1 | 1.0 ± 0.1 | **1.6 ± 0.3** | **1.6 ± 0.3** | 0.6 ± 0.1 | 0.6 ± 0.1 | **0.7 ± 0.1** | **0.8 ± 0.1** |
| Shoot : Root ratio | 2.1 ± 0.6 | 1.6 ± 0.2 | **1.6 ± 0.2** | **1.7 ± 0.9** | 1.9 ± 0.2 | 1.8 ± 0.2 | **2.4 ± 0.5** | **2.0 ± 0.3** | 1.5 ± 0.2 | 1.3 ± 0.1 | **1.3 ± 0.1** | **1.3 ± 0.2** |
| Main tiller dry matter (g) | 0.1 ± 0.0 | 0.1 ± 0.0 | **0.2 ± 0.0** | **0.2 ± 0.0** | 0.2 ± 0.1 | 0.3 ± 0.1 | **0.5 ± 0.1** | **0.4 ± 0.1** | 0.2 ± 0.0 | 0.2 ± 0.0 | **0.2 ± 0.0** | **0.2 ± 0.0** |
| Tillers dry matter (g) | 0.2 ± 0.1 | 0.2 ± 0.0 | **0.4 ± 0.1** | **0.2 ± 0.1** | 0.4 ± 0.1 | 0.4 ± 0.1 | **0.6 ± 0.3** | **0.6 ± 0.3** | 0.2 ± 0.1 | 0.2 ± 0.1 | **0.2 ± 0.1** | **0.2 ± 0.1** |
| Transpiration rate (g day$^{-1}$) | 20.9 ± 2.3 | 23.7 ± 2.0 | **24.0 ± 5.6** | **20.7 ± 5.6** | 34.1 ± 3.2 | 32.3 ± 6.6 | **27.6 ± 7.5** | **26.5 ± 5.1** | 23.3 ± 3.3 | 22.5 ± 2.9 | **18.2 ± 2.7** | **20.6 ± 2.5** |
| Cumulated transpiration (g) | 146.0 ± 81.1 | 171.6 ± 13.9 | **119.2 ± 29.6** | **109.8 ± 25.0** | 333.8 ± 31.1 | 323.0 ± 45.3 | **244.1 ± 49.2** | **246.5 ± 36.1** | 208.4 ± 32.0 | 209.1 ± 27.8 | **126.5 ± 29.5** | **152.6 ± 30.6** |
| WUE (g L$^{-1}$) | 0.9 ± 0.1 | 1.2 ± 0.8 | **1.0 ± 0.31** | **1.7 ± 0.9** | 2.9 ± 1.9 | 3.0 ± 0.8 | **6.4 ± 2.2** | **6.4 ± 3.4** | 2.5 ± 1.0 | 2.5 ± 1.1 | **5.5 ± 2.9** | **4.5 ± 2.3** |
| Mid-drought | | | | | | | | | | | | |
| Time since drought start (day) | 7 | | **11** | | 6 | | **9** | | | | | |
| LER (cm tiller$^{-1}$ day$^{-1}$) | 1.7 ± 0.9 | 0.4 ± 0.3 | **4.6 ± 1.5** | **1.8 ± 0.8** | 2.6 ± 2.2 | 0.2 ± 0.3 | **2.8 ± 1.4** | **0.8 ± 0.6** | | | | |
| Haun index | 7.5 ± 0.5 | 7.4 ± 0.7 | **8.1 ± 0.8** | **7.8 ± 0.5** | 6.6 ± 0.4 | 6.3 ± 0.5 | **6.6 ± 0.3** | **6.4 ± 0.5** | | | | |
| Leaf length (cm) | 12.5 ± 4.4 | 12.4 ± 4.5 | **16.7 ± 8.3** | **16.3 ± 7.0** | 16.1 ± 7.4 | 17.0 ± 7.6 | **20.9 ± 11.7** | **20.2 ± 11.1** | | | | |
| Leaf area (cm²) | 104.6 ± 25.3 | 64.6 ± 10.3 | **228.2 ± 55.7** | **241.6 ± 62.4** | 124.4 ± 52.1 | 107.5 ± 23.4 | **242.1 ± 78.6** | **171.8 ± 24.3** | | | | |
| Specific Leaf Area (SLA, cm² g$^{-1}$) | 197.1 ± 25.0 | 205.1 ± 26.6 | **161.3 ± 49.6** | **138.3 ± 41.0** | 263.6 ± 24.2 | 254.9 ± 51.8 | **154.3 ± 35.8** | **135.2 ± 69.7** | | | | |
| Plant dry matter (g) | 1.1 ± 0.3 | 0.8 ± 0.2 | **3.6 ± 0.9** | **3.2 ± 0.4** | 1.5 ± 0.5 | 1.5 ± 0.2 | **3.4 ± 0.5** | **3.7 ± 1.1** | | | | |
| Shoot : Root ratio | 1.2 ± 0.2 | 1.0 ± 0.5 | **2.1 ± 0.6** | **1.2 ± 0.2** | 1.3 ± 0.3 | 1.1 ± 0.2 | **1.7 ± 0.3** | **1.2 ± 0.2** | | | | |
| Main tiller dry matter (g) | 0.1 ± 0.0 | 0.1 ± 0.0 | **0.3 ± 0.1** | **0.2 ± 0.1** | 0.3 ± 0.1 | 0.3 ± 0.1 | **0.8 ± 0.2** | **0.7 ± 0.2** | | | | |
| Tillers dry matter (g) | 0.4 ± 0.3 | 0.3 ± 0.2 | **2.1 ± 0.9** | **1.5 ± 0.4** | 0.5 ± 0.5 | 0.5 ± 0.2 | **1.3 ± 0.5** | **1.3 ± 1.1** | | | | |
| Transpiration rate (g day$^{-1}$) | 27.0 ± 3.4 | 8.1 ± 3.8 | **55.5 ± 11.7** | **16.0 ± 7.3** | 55.9 ± 7.9 | 6.3 ± 4.2 | **44.2 ± 11.5** | **14.3 ± 4.3** | | | | |
| Cumulated transpiration (g) | 322.7 ± 30.7 | 296.7 ± 19.8 | **573.8 ± 122.9** | **415.3 ± 67.5** | 559.7 ± 64.9 | 422.9 ± 56.2 | **588.5 ± 115.5** | **474.3 ± 91.1** | | | | |
| WUE (g L$^{-1}$) | 3.3 ± 2.0 | 3.1 ± 1.7 | **6.3 ± 4.3** | **7.6 ± 3.4** | 2.6 ± 2.0 | 3.4 ± 1.9 | **5.7 ± 3.2** | **7.7 ± 4.6** | | | | |
| Growth cessation | | | | | | | | | | | | |
| Time since drought start (day) | 11 | | **14** | | 9 | | **12** | | 10 | | **13** | |
| LER (cm tiller$^{-1}$ day$^{-1}$) | 1.8 ± 0.8 | 0.0 ± 0.0 | **2.6 ± 1.0** | **0.0 ± 0.0** | 2.9 ± 0.7 | 0.0 ± 0.1 | **3.3 ± 1.8** | **0.0 ± 0.0** | 2.3 ± 0.5 | 0.1 ± 0.2 | **3.2 ± 1.1** | **0.2 ± 0.2** |
| Haun index | 8.1 ± 0.6 | 7.5 ± 0.7 | **9.0 ± 0.9** | **7.9 ± 0.6** | 6.9 ± 0.6 | 6.3 ± 0.7 | **7.2 ± 0.4** | **6.5 ± 0.5** | 8.3 ± 0.4 | 7.8 ± 0.8 | **9.1 ± 0.3** | **8.0 ± 0.5** |
| Leaf length (cm) | 12.5 ± 4.6 | 12.5 ± 4.5 | **18.8 ± 9.3** | **16.5 ± 7.1** | 17.3 ± 8.0 | 16.9 ± 7.6 | **22.8 ± 11.9** | **20.1 ± 11.2** | 16.5 ± 5.8 | 14.7 ± 4.6 | **17.3 ± 5.1** | **15.6 ± 4.1** |
| Leaf area (cm²) | 127.6 ± 9.2 | 40.4 ± 14.8 | **361.4 ± 85.6** | **89.4 ± 45.9** | 165.0 ± 43.6 | 76.1 ± 27.0 | **285.0 ± 58.3** | **122.1 ± 36.1** | 184.1 ± 30.0 | 74.5 ± 16.5 | **263.5 ± 27.6** | **115.1 ± 16.9** |
| Specific Leaf Area (SLA, cm² g$^{-1}$) | 181.1 ± 13.9 | 148.3 ± 61.8 | **159.5 ± 23.6** | **83.5 ± 7.4** | 250.8 ± 43.6 | 154.9 ± 60.9 | **143.4 ± 44.5** | **77.3 ± 21.4** | 265.6 ± 4.1 | 238.6 ± 55.1 | **227.8 ± 23.3** | **145.6 ± 15.4** |
| Plant dry matter (g) | 1.5 ± 0.3 | 1.0 ± 0.1 | **6.9 ± 1.8** | **3.9 ± 0.7** | 1.9 ± 0.6 | 1.5 ± 0.1 | **4.5 ± 2.0** | **4.1 ± 0.7** | 1.6 ± 0.2 | 1.0 ± 0.1 | **3.6 ± 0.5** | **3.2 ± 0.5** |
| Shoot : Root ratio | 1.1 ± 0.1 | 0.9 ± 0.4 | **1.7 ± 0.2** | **1.0 ± 0.2** | 1.6 ± 0.4 | 1.1 ± 0.5 | **1.8 ± 0.5** | **1.2 ± 0.2** | 1.0 ± 0.2 | 0.6 ± 0.9 | **0.8 ± 0.1** | **0.6 ± 0.1** |
| Main tiller dry matter (g) | 0.2 ± 0.0 | 0.1 ± 0.0 | **0.5 ± 0.2** | **0.2 ± 0.1** | 0.4 ± 0.0 | 0.3 ± 0.1 | **0.7 ± 0.2** | **0.7 ± 0.1** | 0.4 ± 0.4 | 0.2 ± 0.0 | **0.5 ± 0.1** | **0.4 ± 0.1** |
| Tillers dry matter (g) | 0.6 ± 0.4 | 0.3 ± 0.1 | **3.9 ± 1.8** | **1.6 ± 0.6** | 0.8 ± 0.6 | 0.5 ± 0.1 | **2.1 ± 1.5** | **1.5 ± 0.7** | 0.5 ± 0.4 | 0.2 ± 0.9 | **1.1 ± 0.5** | **0.8 ± 0.5** |
| Transpiration rate (g day$^{-1}$) | 33.8 ± 4.1 | 2.3 ± 3.2 | **33.3 ± 14.0** | **0.9 ± 1.3** | 68.1 ± 5.6 | 2.6 ± 4.8 | **62.9 ± 12.1** | **6.7 ± 6.0** | 43.4 ± 7.4 | 4.9 ± 4.0 | **41.5 ± 5.6** | **3.9 ± 2.6** |
| Cumulated transpiration (g) | 541.2 ± 61.4 | 342.7 ± 48.8 | **758.9 ± 140.5** | **412.3 ± 36.7** | 827.1 ± 38.4 | 491.6 ± 20.1 | **779.1 ±81.3** | **523.1 ± 108.6** | 627.3 ± 34.2 | 326.2 ± 0.5 | **582.0 ± 37.0** | **354.7 ± 57.2** |
| WUE (g L$^{-1}$) | 3.3 ± 1.9 | 3.0 ± 2.7 | **9.1 ± 3.5** | **9.9 ± 3.2** | 2.6 ± 0.5 | 3.5 ± 2.4 | **7.6 ± 4.0** | **8.5 ± 5.5** | 3.0 ± 0.5 | 3.0 ± 0.6 | **6.2 ± 2.7** | **8.8 ± 2.8** |
| $gs_w$ (mol m$^{-2}$ s$^{-1}$) | 0.2 ± 0.1 | | **0.06 ± 0.1** | | 0.3 ± 0.1 | | **0.2 ± 0.07** | | 0.27 ± 0.1 | | **0.24 ± 0.1** | |

**Table S6. Mean ± standard deviation data of 10 plants per treatment of water-soluble carbohydrates (% dry matter), osmotic potential (MPa), turgor potential (MPa), water potential of mature leaves and growing zone (MPa), potential gradient of mature leaf and growing zone (MPa), for *Lolium perenne L.*, *Festuca arundinacea* and *Triticum aestivum L.*, at three drought step: drought onset, mid-drought, growth cessation.**

| | RyeGrass | | | | Tall Fescue | | | | Wheat | | | |
|---|---|---|---|---|---|---|---|---|---|---|---|---|
| | 200 ppm | 200 ppm | **800 ppm** | **800 ppm** | 200 ppm | 200 ppm | **800 ppm** | **800 ppm** | 200 ppm | 200 ppm | **800 ppm** | **800 ppm** |
| | Watered | Unwatered | **Watered** | **Unwatered** | Watered | Unwatered | **Watered** | **Unwatered** | Watered | Unwatered | **Watered** | **Unwatered** |
| | Drought onset | | | | | | | | | | | |
| Time since drought start (day) | 0 | | **0** | | 0 | | **0** | | 0 | | **0** | |
| Water-soluble carbohydrates content (%DM) | 5.4 | 6.0 | **15.5** | **13.7** | 11.1 | 11.5 | **32.5** | **24.3** | 11.3 | 13.6 | **17.5** | **16.1** |
| Osmotic potential (MPa) | -0.9 | -0.9 | **-0.9** | **-1.0** | -1.1 | -1.2 | **-0.8** | **-1.1** | -0.6 | -0.6 | **-0.9** | **-0.7** |
| Water potential of Growth Zone (MPa) | -0.9 ± 0.2 | -0.6 ± 0.2 | **-0.6 ± 0.1** | **-0.6 ± 0.3** | -0.6 ± 0.1 | -0.7 ± 0.4 | **-0.7 ± 0.2** | **-0.6 ± 0.5** | -0.5 ± 0.1 | -0.5 ± 0.3 | **-0.5 ± 0.2** | **-0.5 ± 0.1** |
| | Mid-drought | | | | | | | | | | | |
| Time since drought start (day) | 7 | | **11** | | 6 | | **9** | | | | | |
| Water-soluble carbohydrates content (%DM) | 8.5 | 8.6 | **9.4** | **16.1** | 11.3 | 10.6 | **22.0** | **32.6** | | | | |
| Osmotic potential (MPa) | -0.9 | -2.6 | **-0.4** | **-1.0** | -1.0 | -2.1 | **-1.1** | **-1.1** | | | | |
| Water potential of Growth Zone (MPa) | -0.8 ± 0.3 | -1.4 ± 0.4 | **-0.5 ± 0.3** | **-1.0 ± 0.2** | -0.6 ± 0.1 | -1.4 ± 0.3 | **-0.5 ± 0.1** | **-1.0 ± 0.4** | | | | |
| | Growth cessation | | | | | | | | | | | |
| Time since drought start (day) | 11 | | **14** | | 9 | | **12** | | 10 | | **13** | |
| Water-soluble carbohydrates content (%DM) | 8.0 | 9.2 | **21.2** | **14.7** | 6.1 | 8.4 | **22.7** | **31.9** | 6.7 | 7.3 | **13.6** | **37.9** |
| Osmotic potential (MPa) | -1.3 | N.A. | **-1.2** | **-2.1** | -0.9 | -1.6 | **-1.0** | **-2.7** | -0.9 | N.A. | **-0.9** | **-1.8** |
| Water potential of Growth Zone (MPa) | -0.8 ± 0.2 | -1.5 ± 0.8 | **-1.0 ± 0.4** | **-1.6 ± 0.5** | -0.6 ± 0.1 | -1.2 ± 0.5 | **-0.6 ± 0.3** | **-1.9 ± 1.4** | -0.7 ± 0.1 | -1.2 ± 0.5 | **-0.8 ± 0.5** | **-1.0 ± 0.5** |
